# A map of integrated cis-regulatory elements enhances gene regulatory analysis in maize

**DOI:** 10.1101/2024.08.01.606127

**Authors:** Jasper Staut, Nicolás Manosalva Pérez, Andrés Matres Ferrando, Indeewari Dissanayake, Klaas Vandepoele

**Affiliations:** Ghent University, Department of Plant Biotechnology and Bioinformatics, Technologiepark 71, 9052 Ghent, Belgium; VIB-UGent Center for Plant Systems Biology, Technologiepark 71, 9052 Ghent, Belgium; VIB Center for AI & Computational Biology, VIB, Ghent, Belgium

## Abstract

Cis-regulatory elements (CREs) are non-coding DNA sequences that modulate gene expression. Their identification is critical to study the transcriptional regulation of genes controlling key traits that govern plant growth and development. They are also crucial components for the delineation of gene regulatory networks, which represent regulatory interactions between transcription factors (TFs) and target genes. In maize, CREs have been profiled using different computational and experimental methods, but the extent to which these methods complement each other in identifying functional CREs is unclear. Here, we report the data-driven integration of different maize CRE profiling methods to optimize the capture of experimentally-confirmed TF binding sites, resulting in maps of integrated CREs (iCREs) showing increased levels of completeness and precision. We combined the iCREs with a wide diversity of gene expression data under drought conditions to perform motif enrichment and infer drought-specific GRNs. Mining these organ-specific GRNs revealed known and novel candidate regulators of maize drought response, and showed these networks significantly overlap with drought eQTL regulatory interactions. Furthermore, by studying the transposable elements (TEs) overlapping with iCREs, we identified few TE superfamilies displaying typical epigenetic features of regulatory DNA that are potentially involved in wiring specific TF-target gene regulatory interactions. Overall, our study showcases the integration of different omics data sources to generate a high-quality collection of CREs, together with their applicability to better characterize gene regulation in the complex maize genome.

## Introduction

The characterization of functional genomic elements, particularly cis-regulatory elements (CREs), is vital to understand the regulation of complex gene expression programs that coordinate the development and environmental adaptation in plants (Marand *et al*., 2023). CREs contain short and specific non-coding DNA sequences that are bound by transcription factors (TFs) to regulate the expression of genes. These sequences are also termed TF binding sites (TFBSs) or motifs. The comprehensive identification of CREs is key to study gene regulation and identify gene regulatory networks (GRNs), which represent regulatory interactions between TFs and their target genes. GRNs have been widely used to discover the molecular regulators in signaling pathways that regulate plant growth and response to stresses (Chen *et al*., 2018; Clark *et al*., 2021; De Clercq *et al*., 2021; Depuydt *et al*., 2023; Gaudinier *et al*., 2018; Kajala *et al*., 2021; Maher *et al*., 2018; Reynoso *et al*., 2019; Sullivan et al., 2014; Zander et al., 2020; Zhou et al., 2020). In maize, the second most-produced crop worldwide, the characterization of CREs and inference of GRNs is challenging given its large and repeat-rich genome. The maize genome has long intergenic regions, long-range and distal regulatory interactions, and consists of 80% transposable elements (TEs) (Jiao *et al*., 2017; Li *et al*., 2019; Lu *et al*., 2019; Ricci *et al*., 2019). TE proliferation during maize evolution shaped its genome function and structure, and although they are mostly silenced and inactive, they can influence gene expression if they carry CREs upon their transposition in the proximity of genes (Marand *et al*., 2023). Additionally, the insertion of TEs can also disrupt existing CREs or alter the surrounding epigenetic context due to the spreading of repressive chromatin marks (Hirsch and Springer, 2017; Marand *et al*., 2023; Schmitz *et al*., 2022).

Both experimental and computational methods can be employed to identify CREs in a genome-wide manner. Chromatin immunoprecipitation sequencing (ChIP-seq) profiles the *in vivo* binding of one specific TF in a specific context, making it a low-throughput method. The characterization of accessible chromatin regions (ACRs), a TF-independent method to identify putative CREs, can be achieved using various techniques such as assay for transposase accessible chromatin sequencing (ATAC-seq) (Buenrostro *et al*., 2015), DNase I hypersensitive sites sequencing (DNase-seq) (Boyle *et al*., 2008), micrococcal nuclease sequencing (MNase-seq) (Yuan *et al*., 2005), and MNase-defined cistrome-occupancy analysis (MOA-seq) (Savadel *et al*., 2021). These methods aid the identification of CREs because regions depleted of nucleosomes allow TFs to bind to their cognate CREs, hinting at functional regulatory regions in a specific *in vivo* context (Noll, 1974; Oudet *et al*., 1975; Wang *et al*., 2011; Weintraub and Groudine, 1976). DNA methylation is also involved in gene regulation: hypomethylated regions tend to overlap with ACRs (Eli *et al*., 2016; Oka *et al*., 2017; Ricci *et al*., 2019), while hypermethylation is usually observed in silenced genomic regions (Soppe *et al*., 2002). Unmethylated regions (UMRs), profiled with deep whole-genome bisulfite sequencing, were shown to be located nearby genes expressed and to overlap with ACRs in tissues different from the one profiled. Furthermore, DNA methylation patterns in plants show low variability during the vegetative development (Eichten *et al*., 2013; Kawakatsu *et al*., 2016) and in response to environmental changes (Crisp *et al*., 2017; Eichten and Springer, 2015; Ganguly *et al*., 2017). Thus, UMRs show potential to identify CREs that are functional in different organs/tissues, developmental stages, or environments, allowing to profile a more complete set of regulatory elements compared to ACRs (Crisp *et al*., 2020).

Computational methods that characterize putative CREs focus mainly on profiling genome conservation to identify conserved non-coding sequences (CNSs). CREs tend to show increased sequence conservation as a result of purifying selection due to their role in gene regulation (Cooper and Brown, 2008; Haudry *et al*., 2013; Kimura, 1991). FunTFBS (Tian *et al*., 2020), *msa_pipeline* (Wu *et al*., 2022), BLSSpeller (De Witte *et al*., 2015), or the method developed by (Song *et al*., 2021) are examples of approaches that have been employed to identify CNSs in plants, including maize. FunTFBS and *msa_pipeline* assess genome-wide conservation using chained pairwise genome alignments together with evolutionary models to compute conservation scores and identify conserved elements. BLSSpeller uses k-mers to assess the conservation of DNA motifs in the promoter sequences of orthologous and paralogous genes, taking relative phylogenetic distances into account. The method of (Song *et al*., 2021) uses pairwise alignments to identify extended seeds, representing locally conserved sequences, as CNSs. The Conservatory project (Hendelman *et al*., 2021) (https://conservatorycns.com/) uses cis-regulatory sequences and gene-based microsynteny to derive multiple alignments of upstream and downstream regulatory sequences in closely related species. It then reconstructs the ancestral sequences for each CNS and searches for that in syntenic regions of distant genomes.

In contrast to TF ChIP-seq profiling, profiling chromatin accessibility, DNA methylation or genome conservation provides a more general, context-independent way to delineate a complete map of CREs. While accurate CRE identification aids the characterization of regulatory DNA flanking individual genes, it also enables the development of motif-based tools that predict GRNs, supporting gene regulation analysis (Ding *et al*., 2021; Ferrari *et al*., 2022; Kulkarni *et al*., 2018; Manosalva Pérez *et al*., 2024; Wilkins *et al*., 2016). Despite the availability of diverse methods that characterize putative CREs in plants, it is currently unclear which one is most suited to identify TFBSs and regulatory interactions. Here we present an evaluation of five CNS detection methods in maize, and their comparison and integration with UMRs and a compendium of ACRs. We benchmarked these three CRE sources to assess their complementarity and generate a map of integrated CREs (iCREs). We demonstrated the power of the iCREs in identifying functional TFBSs by inferring iCREs-based GRNs that are specific for maize drought response across different organs. These GRNs allowed identifying diverse drought TFs and shed light on the downstream processes they control, as well as their organ specificity. Additionally, we found specific TE superfamilies that are enriched in iCREs, display chromatin signatures of regulatory DNA, are overrepresented for specific TFBSs, and influence gene expression.

## Results

### Complementarity between CREs identified using computational and experimental approaches

Different CRE-profiling methods exist, but an evaluation of their complementarity at predicting functional TFBSs is missing in plants. We evaluated five CNS detection methods that have been applied in plants (BLSSpeller, *msa_pipeline*, funTFBS, a method developed by (Song *et al*., 2021), and the method of the Conservatory project), along with two experimental methods used to profile epigenetic landscapes (ACRs and UMRs), as candidate approaches to detect putative CREs. In order to evaluate and compare their ability to detect regulatory DNA, in particular TFBSs, we used a ChIP-seq gold standard that comprises binding events of 104 TFs in maize mesophyll cells (Tu *et al*., 2020). As chromatin accessibility indicates regions that can be bound by TFs, we considered the ACR and UMR datasets included in our benchmark as two alternative, silver standards to help us assess the robustness of our findings. To obtain a compendium of ACRs, we merged 13 high-quality datasets across different tissues (leaf, root, ear, tassel, husk, inner stem, above ground seedling, and axillary bud; see Methods; Figure S1). As a control, we included repetitive DNA and TEs in our benchmark, which are known to be depleted of CREs (Marand *et al*., 2023).

We found that predictions of all the tested methods were significantly enriched (p-value < 0.05) for the putative binding sites of the ChIP-seq data, while the repeats/TEs were depleted (Table S1), demonstrating the ability of each method to identify TF binding events. Next, we computed the precision, recall, and F1 of all methods by overlapping them (a base pair intersection) with the ChIP-seq dataset as ground truth. Precision is a metric for the correctness of the predictions, while recall reports the completeness (i.e., how many regions in the gold standard were identified). F1 is the harmonic mean between precision and recall, balancing correctness and completeness. We observed notable differences in performance between methods, in particular among the CNS detection methods (Figure 1A; Table S1). The method by (Song *et al*., 2021) strongly outperformed other CNS detection methods in recall (62.9%) and was in precision (13.3%) only surpassed by Conservatory (22.0%). To assess the robustness of these trends, we used the compendium of ACR datasets and the UMR dataset as alternative silver standards and observed similar patterns when predicting ACRs or UMRs (Figure S2). Taken together, we find the method by (Song *et al*., 2021) to be the best performing CNS method as quantified by F1 (22.0% for the ChIP-seq gold standard). However, both ACRs and UMRs show a higher F1 than the best-performing CNS detection method (Figure 1A; Table S1). Among these two, UMRs are the best at predicting the TF binding events of the gold standard (1.7% higher precision and 15.9% higher recall than ACRs). The evaluation with the ACR silver standard (where ACRs are excluded from the evaluation) further confirms that UMRs outperform all CNS detection methods (Figure S2A).

**Figure 1.**
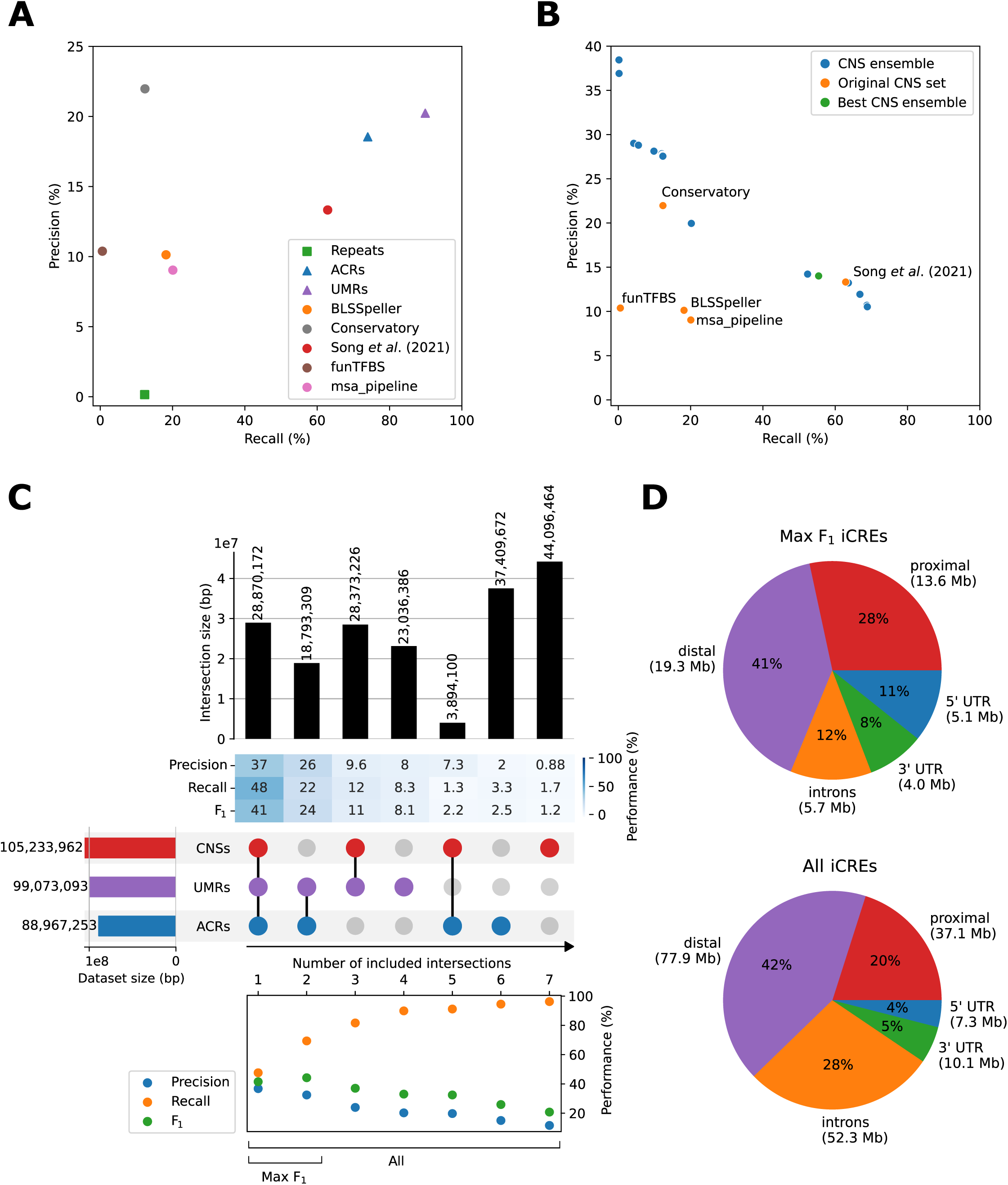
Benchmark, complementarity assessment, and integration of CNS detection and experimental CRE-profiling methods. (A) Precision-recall plot of individual CNS detection and experimental CRE-profiling methods, using a ChIP-seq dataset as gold standard. CNS detection methods are plotted as circles, experimental CRE-profiling methods as triangles, and control datasets as squares. (B) Precision-recall plot (using ChIP-seq as gold standard) of individual CNS detection methods and different ensembles that were constructed by combining the intersections of these methods. The ensemble with the best F1 is shown in green. (C) UpSet plot of the ACRs, UMRs and CNSs, with performance metrics of the subsets. The UpSet plot shows the sizes of the full datasets on the left and the sizes of the unique subsets on the top. Precision, recall and F1 are calculated using a ChIP-seq gold standard for each of the individual subsets and are shown in a heatmap below the size bars. The UpSet plot is sorted on the precision of the subsets, so that the subset with the most correct predictions is shown on the left. Precision, recall and F1 are also calculated for different ensemble sets, created by starting from the subset with the highest precision (most left) and progressively adding individual subsets with the next best precision, until all regions are added together (most right). The combined subsets that result in the “max F1 iCREs” and “all iCREs” are labeled “max F1” and “all” respectively. (D) The genomic feature distribution of the “max F1 iCREs” and “all iCREs”. The total coverage in each genomic feature is given in megabases (Mb).

Based on our results indicating that both the CNS detection and experimental datasets identify functional regulatory DNA, we next assessed their complementarity, starting with the CNS datasets. We quantified the overlap (in base pairs) of the five CNS sets and visualized them in an UpSet plot, with each column showing a unique (i.e., not overlapping with another column) subset of genomic regions (Figure S3). In addition, we calculated the precision, recall, and F1 of each unique subset shown in the UpSet plot. As expected, the intersection of all five CNS detection methods has the highest precision. Next, we evaluated if an ensemble of different CNS detection methods can outperform the best performing individual method. Therefore, we took the unique subset of genomic regions with the highest precision and subsequently added the subset with the next best precision until all subsets were included. Evaluating precision, recall and F1 with every added subset (ensemble F1) shows that an ensemble approach can yield a set of CNSs with a much higher precision (47.0%) than the best individual CNS set (22.0%), albeit at the cost of a reduced recall (Figure 1B; Figure S3). However, when comparing the CNSs of (Song *et al*., 2021) with an ensemble of similar recall, both have similar performances (the ensemble’s recall is 0.27% higher and its precision is 0.00001% higher). We observed that the best CNS ensemble only shows a marginal improvement in F1 (0.38% higher) over the best individual method, demonstrating that the ensemble is not able to substantially outperform the method of (Song *et al*., 2021). Thus, for simplicity, we did not continue with a CNS ensemble approach. Instead, we used the CNSs of (Song *et al*., 2021) to assess the complementarity between CNSs, ACRs and UMRs for the purpose of integrating them into a map of putative CREs.

To integrate CNSs, ACRs and UMRs in a way that maximizes the agreement with the ChIP-seq gold standard and thereby optimally captures functional CREs, we performed a complementarity analysis, in the same way as with the CNS detection methods. Unique subsets of genomic regions were defined by overlapping and subtracting the different methods and then calculating the performance for each unique subset (Figure 1C). Each of the three methods contains a substantial fraction of unique regions. Regions unique to UMRs show a better agreement with ChIP-seq-confirmed bindings sites than unique ACRs or CNSs, as well as regions where ACRs and CNSs overlap but do not coincide with a UMR. In addition, we observe that combining methylation data with other methods (CNSs or ACRs) can further improve precision (up to 37%, compared to 20% when considering all UMRs; Figure 1C). This result reveals that genome conservation, chromatin accessibility and DNA methylation contain complementary information to predict CREs. Finally, we integrated the CNSs, ACRs and UMRs into a set of “integrated CREs” or iCREs. As described before, we combined the unique subsets shown in the UpSet into ensembles by progressively adding the subset with the next best precision, and selected the ensemble with the highest F1 (Figure 1C) using the ChIP-seq gold standard. This ensemble, referred to as the “max F1 iCREs”, is equivalent to the intersection between the UMRs and the ACRs. We then constructed a second ensemble consisting of the union of CNSs, ACRs and UMRs that contains the regions of all three methods, which will be referred to as the “all iCREs”. Figure 1D indicates the distribution of the ensemble sets in different genomic regions and/or features. As expected, the “max F1 iCREs” have lower genome coverage (48 Mb) and are more often located in a UTR or proximal region (19% UTR and 28% proximal), while “all iCREs” cover more of the genome (184 Mb) and have a lower proportion in these regions (9% UTR and 20% proximal). In summary, we show that DNA methylation has a high level of agreement with ChIP-seq-confirmed TFBSs and that it can be complemented with chromatin accessibility and/or genome conservation information. This complementarity allows us to generate a high confidence (max F1 iCREs) and a more comprehensive (all iCREs) map of putative maize CREs.

### Inference of drought-responsive gene regulatory networks using the iCREs and an organ-specific gene expression atlas

iCREs represent a comprehensive collection of maize regulatory regions, having the potential to improve GRN inference. Therefore, we implemented a framework to perform iCREs-based motif enrichment and GRN inference starting from a set of functionally-related genes, using the MINI-AC tool (Manosalva Pérez *et al*., 2024) (Figure 2A).

**Figure 2.**
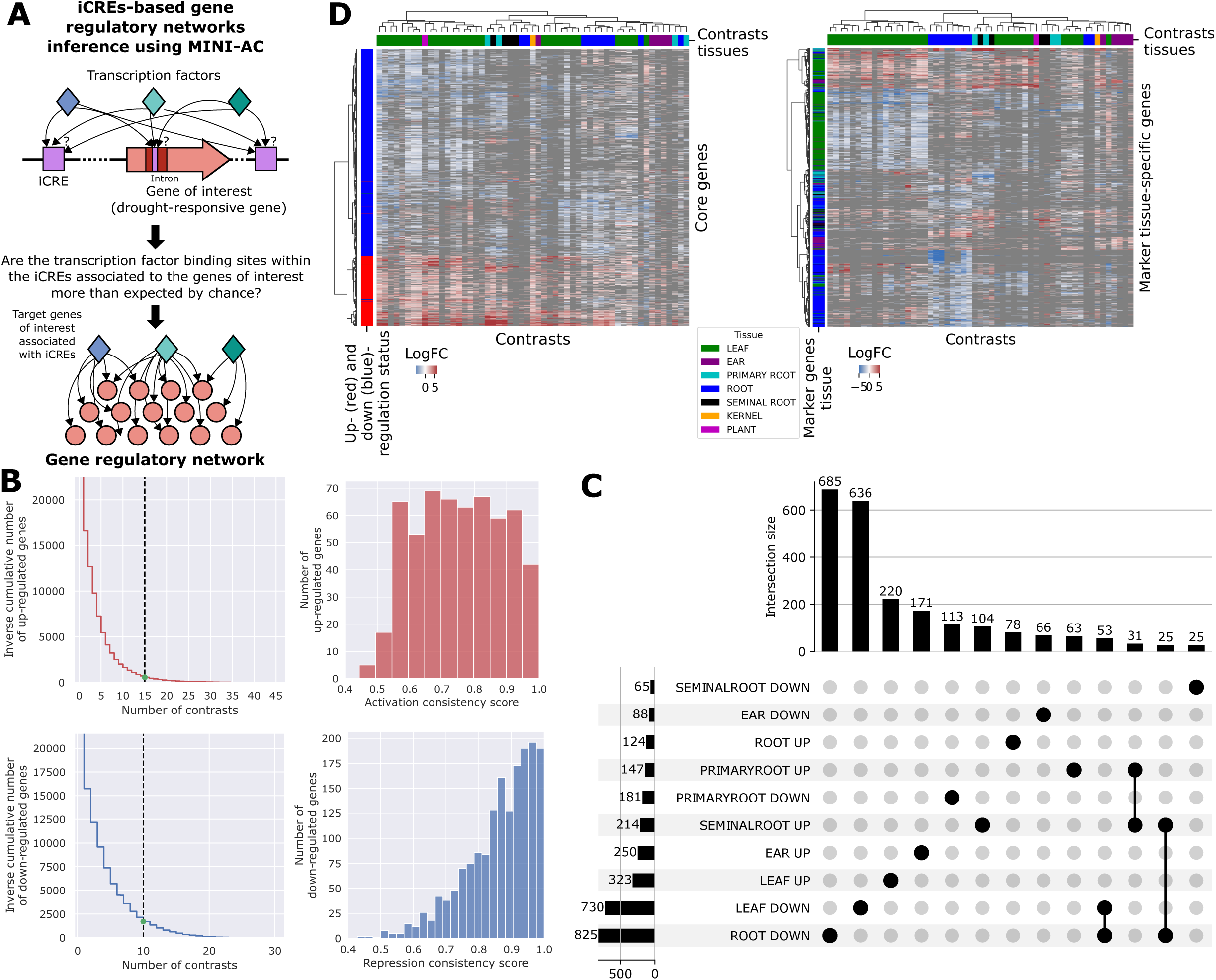
iCREs-based gene regulatory network (GRN) prediction and selection of drought-responsive gene sets. (A) Overview of the iCREs-based GRN inference framework, where the iCREs associated to a set of functionally-related or co-expressed genes are submitted to MINI-AC to perform motif enrichment and GRN inference. (B) On the left, there is the inverse cumulative distribution of the number of up- and down-regulated genes (y-axis) that are common in a specific number of contrasts (x-axis). On the right, there is the activation and repression consistency score distribution of the “core” drought-responsive genes. (C) UpSet plot showing the number of common and shared drought-responsive genes per tissue, sorted by the size of the intersection set. For visualization purposes, only intersection sets with 25 or more genes are shown. (D) Clustered heatmaps showing the log-transformed fold change of gene expression in the “core” (left) and “marker tissue-specific” (right) drought-responsive gene sets.

The frequency and severity of droughts is expected to increase given the current climate emergency, ultimately impacting the yield of maize. To find candidate drought regulators and understand the processes they control, we first constructed a gene expression atlas by selecting and curating 20 maize drought RNA-seq studies to obtain sets of drought-responsive differentially expressed genes (DEGs; Table S2, Figure S4, Materials and Methods). These studies include samples of six different organs: leaf, root, seminal root, primary root, kernel, and ear. Next, we identified “marker” genes that respond to drought exclusively in each of these organs, as well as general or “core” drought-responsive genes that consistently respond to drought across various experimental conditions, samples, and genotypes.

The “core” drought-responsive genes were identified by counting the number of contrasts (the comparison between a control and drought sample) in which each drought-responsive gene is detected as differentially expressed. Next, we selected the genes up- or down-regulated in at least one third of the maximum number of contrasts that share at least one DEG. This yielded 568 “core UP” genes in 15 contrasts and 1702 “core DOWN” genes in 10 contrasts (Figure 2B). Given the diverse selection of studies, there were genes showing repression (i.e. down-regulation) in some contrasts and activation (i.e. up-regulation) in others. Therefore, we quantified the activation or repression consistency of the “core” genes as the proportion of contrasts in which a gene is up- or down-regulated out of all contrasts where it is differentially expressed (see Methods). While we overall observed high consistency scores, they were higher (p-value < 2.2 x 10-16 using a one-sided Mann-Whitney U test) for the down-regulated core genes (median 0.88) than for the up-regulated ones (median 0.75). This trend likely stems from the leaf samples, as this is the organ with most contrasts and the only one replicating low activation consistency scores (Figure 2B and S5). This difference in consistency scores might be due to the drought-induced repression of numerous photosynthetic genes, resulting in higher repression consistency scores (Berrío *et al*., 2022).

The organ-specific drought-responsive genes were defined as the DEGs in more than half of the contrasts of that organ (Figure 2C; kernel was discarded due to the large number of DEGs and having only one contrast). We evaluated the overlap between these organ-specific activated and repressed genes to define unique or “marker” drought-responsive genes. We found that in most organs, the majority of the specific DEGs are unique (Figure 2C and S6). For instance, leaf and root have 87% and 83% of unique repressed DEGs, respectively. Clustering the “core” drought-responsive genes based on their expression log fold-change confirmed a ubiquitous drought response (repression and activation across many contrasts) while clustering the “marker” drought-responsive genes supported their specificity (Figure 2D).

Next, for each drought-responsive gene set, we identified their associated iCREs and performed motif enrichment and GRN inference using MINI-AC (GRNs available as dataset S1; GRN summary statistics in Table S3). The construction of the GRN is done by finding motifs overrepresented in the iCREs associated to a gene set and linking them to the TFs that bind those motifs (MINI-AC normally does this with a set of ACRs). We evaluated if there was an advantage in performing motif enrichment using iCREs as input (iCREs-based enrichment) compared to using the adjacent non-coding regions and introns of the drought-responsive genes (default promoter-based enrichment, see Materials and Methods of iCREs-based GRN inference). Therefore, we compared the performance of these two approaches by measuring how well MINI-AC’s enrichment analysis predicts the motifs bound by a gold standard set of TFs, which are the differentially expressed TFs in each drought-responsive gene set. In other words, we evaluate if the motifs associated to iCREs of drought-responsive DEGs are significantly enriched for drought-responsive TFs. Additionally, we also made a comparison of the “all iCREs” and “max F1 iCREs” sets (Figure 3A; Table S4). Our results showed that the iCREs-based motif enrichment outperforms the promoter-based enrichment in 9 out of the 12 gene sets for the two metrics tested (F1 and the area under the precision recall curve or AUPRC, that measures the performance of the top-scoring predictions; See Materials and Methods of iCREs-based GRN inference). The iCREs-based approach achieves a median F1 10% higher and a median AUPRC 6% higher than the promoter-based approach. Although the “maxF1” and “all” iCREs showed similar performances, the latter was superior for most of the drought-responsive gene sets, with a median 0.049 AUPRC and 0.073 F1, and a median 0.065 AUPRC and 0.1 F1, respectively.

**Figure 3.**
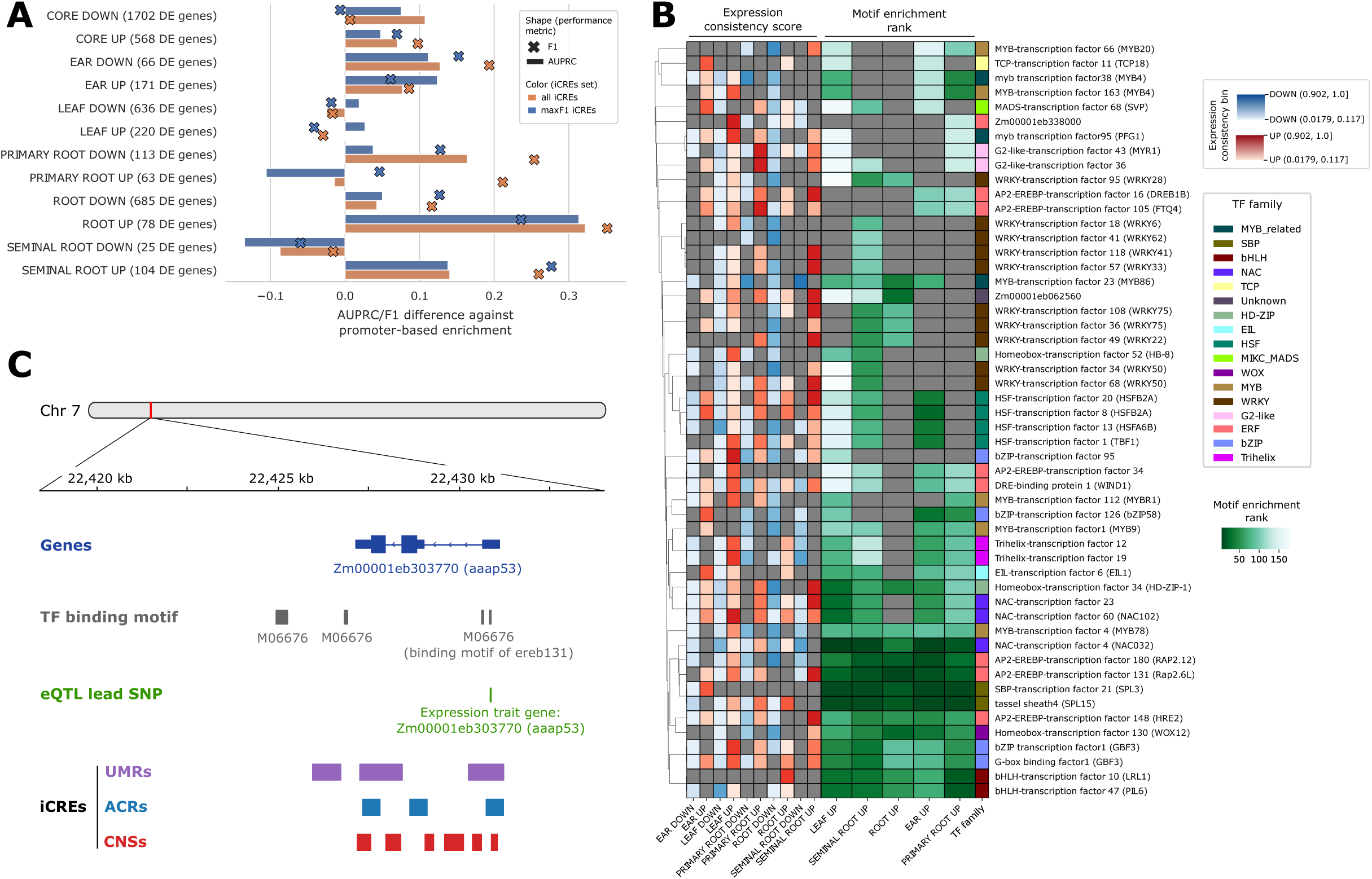
Benchmark and analysis of iCREs-based and drought-responsive gene regulatory networks (GRNs). (A) Comparison of the performance of the “promoter-based” and “iCREs-based” GRN inference frameworks, using two different iCREs sets (maxF1 and all). The two performance metrics (F1 and area under the precision recall curve) measure how well each method predicts differentially expressed (DE) regulators given a set of tissue-specific drought-responsive genes (y-axis). The “promoter-based” results are considered the baseline to which the difference in performance is shown (x-axis). (B) Clustered heatmap showing the motif enrichment rank of different DE transcription factors (TFs) (y-axis) given sets of iCREs that are associated with tissue-specific (x-axis) up-regulated genes in response to drought. The row annotations correspond to the activation and repression consistency score bins and the TF family. For visualization purposes, only TFs with an expression consistency score higher than 0.513 are shown. (C) Visualization of the predicted regulation of aaap53 by TF ereb131. The “Genes” track shows genes within the genome. Exons are shown as rectangles, of which those representing coding sequence have greater height, while introns are shown as lines. The “TF binding motif” track shows the location of the TF ereb131 binding site M06676. The “eQTL lead SNP” shows the location of eQTL lead SNPs, linked to the expression of the trait gene annotated under the SNP. The “iCREs” track shows the location of UMR, ACR and CNS region, included in the iCRE dataset.

To further assess the advantages of using iCREs for GRN inference and the validity of our drought-responsive GRNs, we obtained a set of publicly available drought-specific eQTLs, linking single nucleotide polymorphism (SNP) variants to expression differences of a trait gene (Liu *et al*., 2020) (Materials and Methods). Using the lead SNPs of these drought-specific eQTLs (n=19,154), we find that within the non-coding genome, there is a 22-fold enrichment of the SNPs for the “all iCREs” (p-value < 0.001), showing strong preference of these SNPs to be located in an iCRE. Furthermore, while the “all iCREs” set reduces the number of genome-wide motif occurrences of MINI-AC’s motif mapping by 90%, it only reduces the number of drought eQTL SNPs by 25%. This validates the assumption that mostly false positive motif matches are removed by applying the iCREs, thereby improving the resulting GRNs.

To evaluate the drought-responsive GRNs against the experimental drought eQTL data, we constructed an eQTL-based GRN. We took the expression trait genes as target genes, and linking them to a TF if the lead SNP is located in a binding motif of that TF. We then compared the number of overlapping edges between the eQTL GRN and the real MINI-AC core drought GRN (summary statistics in Table S3), against a set of background GRNs constructed by shuffling the edges of the MINI-AC GRN. This analysis shows that the promoter-based MINI-AC GRN has an 8% higher overlap (35,870 edges) with the eQTL GRN than the median background overlap (33,145 edges), while this increased to 20% (13,491 real, 11,292 background) for the “all iCREs” and 31% (8,287 real, 6,382 background) for the “max F1 iCREs” (p-value of all overlaps < 0.001). This further confirms that the reduction of false positive motif matches using the iCREs improves the quality of our drought GRNs.

Next, we analyzed the drought-responsive GRNs using “all iCREs” and the “marker” gene sets, representing organ-specific drought DEGs. Using MINI-AC, motifs were scored based on how overrepresented they are in the iCREs of the DEGs to compute a motif enrichment rank. Comparing the ranks along with the activation and repression consistency in each organ allowed us to pinpoint specific drought stress regulators (Figure 3B and S7). We observed high expression and motif enrichment specificity for WRKY TFs in root cell types, especially in seminal root (WKRY57, WRKY118, WKRY108, and WKRY49). Several WRKY TFs have been characterized for their role in maize drought response (Wang *et al*., 2018; Zhao *et al*., 2021) and have been associated with an increase of lateral roots (Gulzar *et al*., 2021). In leaf, we observed high expression specificity of Trihelix TFs (Trihelix19 and 12) as well as a low (but not specific) motif enrichment rank. The Arabidopsis ortholog of these TFs, GT1, has been characterized for its role in the repression of cell growth (Breuer *et al*., 2012; Caro *et al*., 2012) and the regulation of drought tolerance (Yoo *et al*., 2010), which is in line with the growth arrest that occurs under drought stress (Dubois and Inzé, 2020). bZIP95 in leaf also showed both high expression and motif enrichment specificity, hinting at an important role in the drought response of leaf (Figure 3B). Focusing on leaf, we found an expected repression of photosynthesis- and carbon metabolism-related genes under the control of repressed TFs (Figure S8; Dataset S2). Among these, we find *Arabidopsis thaliana* orthologs (PIL6, MYBH or CRF5) that have been characterized in light signaling and cell expansion (Fujimori *et al*., 2004; Lu *et al*., 2014; Raines *et al*., 2016). Conversely, we found that the aforementioned Trihelix19, Trihelix12 and bZIP95 are controlling genes involved in response to abscisic acid and water deprivation, further supporting their potential role in maize leaf drought response. Finally, in Figure 3C, we visualize an example locus for one of the predicted drought regulators shown in Figure 3B. The TF ereb131 is predicted by MINI-AC as a regulator of aaap53, a target gene showing DE under drought stress. Illustrated in Figure 3C, there is a binding motif for ereb131 in the UTR of its putative target gene, which falls within an iCRE (here both UMR, ACR and CNS). Furthermore, we find that the lead SNP of an eQTL for the expression of aaap53 is located within this binding motif, providing experimental support for this interaction. The ereb131 TF also has an *A. thaliana* ortholog (RAP2.6L), which has been shown to play a role in drought response (Krishnaswamy *et al*., 2011), making this maize TF an interesting candidate for further study. Taken together, these results illustrate the potential of employing the iCREs to infer context-specific GRNs and identify candidate regulators.

### iCREs-guided characterization of the regulatory role of different types of transposable elements

While numerous examples of TEs influencing gene expression have been reported in plants (Deneweth *et al*., 2022; Hirsch and Springer, 2017; Noshay *et al*., 2021), the role of specific types of TEs contributing to regulatory elements is less clear. Therefore, we identified a subset of TEs fully contained within iCREs and examined their potential regulatory role based on the superfamily they belong to. In the maize genome, different TE superfamilies and types of repetitive elements are classified according to their sequence similarity and transposition specificity (Wicker *et al*., 2007) (see Materials and Methods of TE analysis). We assessed whether specific TE superfamilies/repetitive elements were significantly overrepresented in the iCRES. While the full set of TEs is depleted in the iCREs, the superfamilies Tc1/Mariner, PIF/Harbinger, and hAT that belong to the TE order of terminal inverted repeats (TIR), show a high enrichment in iCREs (enrichment folds of 3.5, 2.9, and 2.3, respectively; Figure 4A). Additionally, the helitron superfamily and two superfamilies belonging to the long interspersed nuclear element (LINE) TE order also showed a 1.4- to 2.2-fold enrichment. Using the non-coding and non-TE genomic space as control regions confirms that the iCREs are depleted from TEs, except for these 6 superfamilies. Additionally, a gene ontology (GO) enrichment analysis revealed that the TEs within iCREs belonging to the TIR superfamily are the most significantly associated with specific groups of functionally-related genes (Figure 4B). For example, we found that the responses to (a)biotic stresses such as salt, cold, bacteria, and fungus are among the most enriched GO terms (q-values between 5.5 x 10^-3^ and 2.4 x 10^-8^ and enrichment folds between 1.3 and 1.4), along with developmental functions like regulation of flower development and chloroplast organization. While other TE superfamilies of the LTR and LINE orders also yielded GO enrichment among their associated genes (mostly related with carbohydrate metabolism; Table S5), Figure 4B depicts only the terms with most significant enrichment (q-value < 0.008 with >100 genes in that category).

**Figure 4.**
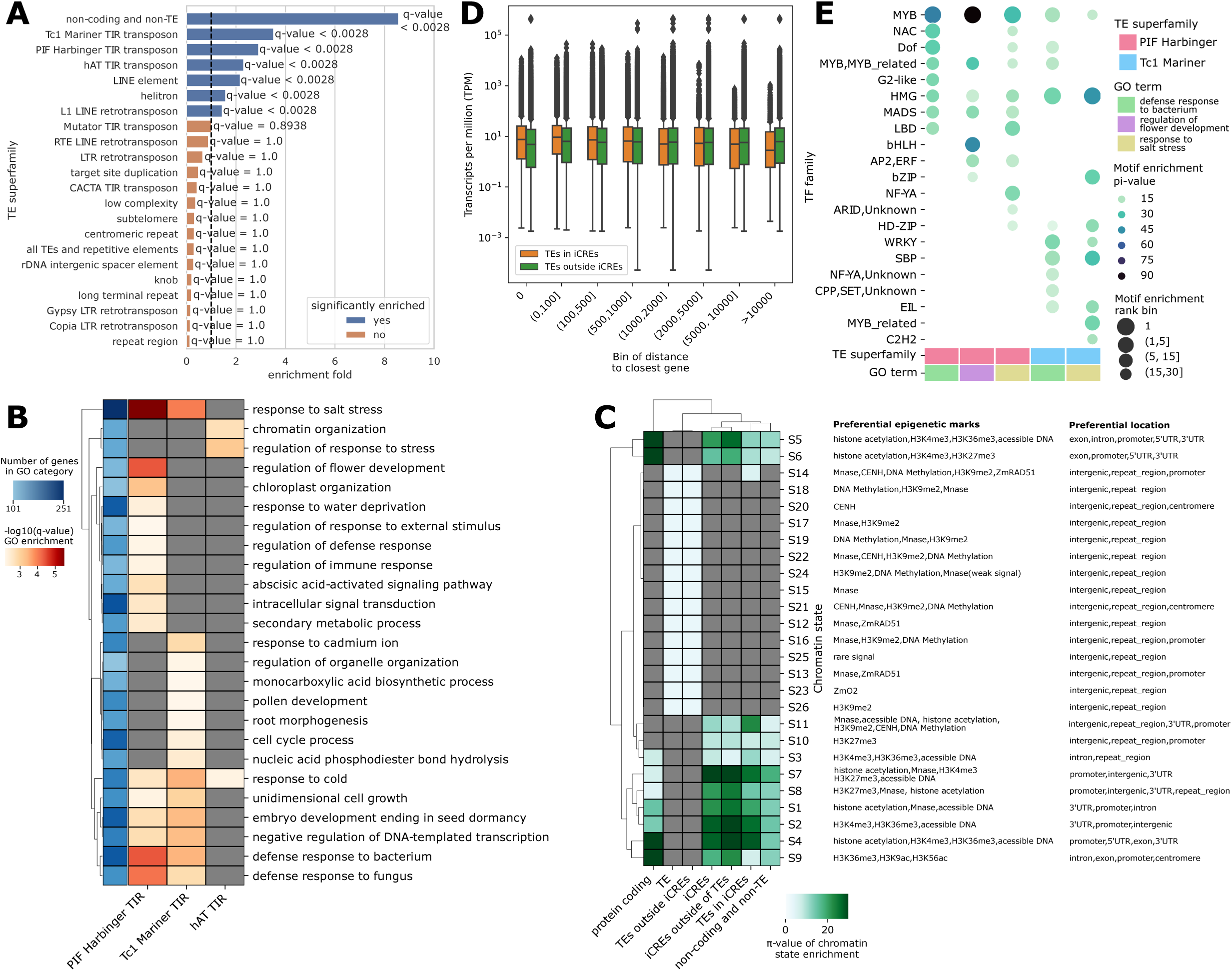
Analysis of the regulatory role of transposable elements (TEs) in maize using iCREs. (A) Bar plot that shows the enrichment fold (x-axis) and q-value of TE superfamilies and repetitive element types (y-axis) in the iCREs. (B) Clustered heatmap displaying the −log10(q-value) of gene ontology (GO; y-axis) enrichment among the genes associated with TEs within iCREs of three tandem inverted repeats (TIR) superfamilies (x-axis). The row annotations indicate the number of TE-associated genes in the corresponding GO category. The heatmap was filtered for TF-GO pairs with a q-value < 0.0008 with at least 100 TE-associated genes within that GO category. (C) Chromatin states (CS; y-axis) enrichment of different sets of genomic regions (x-axis), including all TEs and repetitive elements, iCREs, and TEs within and outside iCREs. The tables adjacent to the x-axis indicate the chromatin marks and genomic locations characteristic of each CS. (D) Box plot displaying the expression level distribution in transcripts per million (y-axis) of the genes associated with TEs within iCREs and outside iCREs, binned per distance to the corresponding genes. (E) Bubble plot showing the motif enrichment rank and π-value (−log10(p-value) × enrichment fold) of different transcription factor (TF) families on the TEs within iCREs that are associated to genes annotated with “defense response to bacterium”, “regulation of flower development”, and “response to salt stress”, and that belong to the superfamilies Tc1/Mariner and PIF/Harbinger. Only the families with at least one motif with motif enrichment rank of 30 and less are shown.

Next, to investigate if the epigenetic profile of the TEs within iCREs is different from the one of the TEs outside iCREs, we compared their overlap with different chromatin states. Chromatin states are sets of genomic regions with specific combinations of epigenetic features that include histone marks, histone variants, DNA methylation, ACRs, and binding sites of chromatin-associated factors (Liu *et al*., 2018). We observed that TEs within iCREs share a similar chromatin state enrichment pattern as the iCREs, which implies that they tend to be in promoter, intronic, UTR, and intergenic regions (Figure 4C). They also display the opposite pattern as the full set of TEs and the TEs outside iCREs, that are enriched in chromatin states associated with repetitive regions showing DNA methylation. Furthermore, we explored the chromatin state enrichment pattern per TE superfamily and repetitive element type, in, and outside iCREs. We found that the helitron and TIR TEs within iCREs, in particular Tc1/Mariner, CACTA, PIF/Harbinger, Mutator, and hAT have the most similar chromatin state enrichment pattern to the iCREs (Figure S9), suggesting they represent functional regulatory DNA.

Besides examining the epigenetic profile of the TEs within iCREs, we also studied expression levels of nearby genes. We first evaluated whether the distribution of the distance to the closest gene was different for the TEs that overlap with iCREs compared to those that do not. We found that TEs are generally closer to genes when they are within iCREs than when they are outside (Figure S10; median distance in iCREs is 7,790 bp vs. 24,705 bp outside iCREs; p-value < 2.2 x 10^-16^ using a one-sided Mann-Whitney U test). Considering this, we used the gene expression atlas of maize drought to evaluate differences in the expression of genes associated with TEs within iCREs, outside iCREs, or genes associated with both. This analysis revealed that in both control and drought samples, the median expression of genes associated with TEs in iCREs is significantly higher than those outside iCREs at distances of less than 2000 bps from the genes or when they are inside the gene body, meaning TEs within introns and UTRs (median TPMs 7.5-8.1 in iCREs vs. 4.8-5.2 outside iCREs at distance 0 from genes; p-value<0.0001 using a one-sided Mann-Whitney U test; Figure 4D; see statistics in Table S6). This trend is also noticeable for the expression values in the control and drought samples when only considering the genes associated with the Tc1/Mariner and PIF/Harbinger superfamilies that display enrichment in chromatin states typical of regulatory DNA (Table S6).

To further explore the regulatory function of these TEs within iCREs, we utilized a public STARR-seq dataset (Ricci *et al*., 2019), quantifying the enhancer activity of maize genomic DNA fragments. We found that the Tc1/Mariner, CACTA, PIF/Harbinger, Mutator, and hAT superfamilies showed significantly (p-value < 2.2 x 10^-16^) increased enhancer activity within iCREs compared to outside iCREs (Table S7). For example, for the CACTA TIR superfamily has a median enhancer activity score of 0.205 in iCREs, but a median score of 0 outside iCREs. This shows that the TEs within iCREs of those TE superfamilies are not only associated with epigenetic marks of regulatory DNA, but also have increased regulatory activity *in vivo*.

Finally, we employed a motif analysis on the PIF/Harbinger and Tc1/Mariner TEs fully overlapping with iCREs, and whose associated genes were functionally annotated with “response to salt stress”, “defense response to bacterium”, or “regulation of flower development” (most enriched GO categories; Figure 4E). We found that the MYB TFs and the high mobility group (HMG) TFs were enriched in all the different test sets. The MYBs were especially enriched in the PIF/Harbinger TEs associated with genes involved in flower development, and the HMG in the Tc1/Mariner TEs associated with genes involved in response to salt stress. Additionally, in the latter, the SBP family also showed high enrichment. The bHLH TF family was highly enriched in the PIF/Harbinger TEs in the context of flower development. In the Tc1/Mariner TEs associated with genes involved in response to bacteria, there was a high enrichment of WRKY TFs, which is expected for this TF family, although this was not the case for the PIF/Harbinger TEs (Chen *et al*., 2019). Next, we investigated whether TF ChIP-seq *in vivo*-profiled binding sites (Tu *et al*., 2020) of the aforementioned TF families were overrepresented within the PIF Harbinger and Tc1-Mariner TE superfamilies (Table S8). We found that the MYB family had an enrichment π-value (see Materials and Methods of TE analysis) of 35.5 in the PIF Harbinger TEs in iCREs, compared to 6.9 outside iCREs. A similar trend was also found for the TF family bHLH (31.5 inside iCREs vs. 6.5 outside). For the Tc1 Mariner superfamily, we found that the MYB and WRKY families had a π-value of 22.9 and 19.8 in iCREs, compared to 4 and 3.9 outside iCREs, respectively. This observation reveals that specific TE superfamilies part of iCREs harbor functional binding sites bound by specific TFs.

Altogether, these results highlight the potential regulatory role of specific TE superfamilies in maize. The presence of overrepresented TFBSs in these TE superfamilies, as well as the prevalence of specific biological processes in the associated genes, suggests that the wiring of these TF families, linking TFs to their target genes, is (partly or completely) mediated by specific TEs.

## Discussion

Integrating datasets from various regulatory DNA profiling methods in plants holds significant potential for enhancing our understanding of gene regulation. CRE-profiling methods such as ChIP-seq can detect direct TF binding, but are relatively low-throughput. Profiling chromatin accessibility, DNA methylation or genome conservation offers a more straightforward way of generating a complete map of CREs. Although previous studies have focused on the agreement between ChIP-seq, CNSs, ACRs and UMRs, their complementarity and potential for integration was largely unexplored (Crisp *et al*., 2020; Marand *et al*., 2023; Ricci *et al*., 2019; Song *et al*., 2021). We show that regions supported by multiple methods are more likely to be confirmed by ChIP-seq, and that each method contains a substantial amount of unique regions, indicating complementarity. While genome conservation provides increased confidence in a region’s regulatory role, our benchmark demonstrates that it is less reliable than profiling methylation or accessibility. Indeed, CNSs are prone to false positives, such as conserved non-coding regulatory RNA, and false negatives, e.g., when a TFBS has been recently introduced in evolutionary time, thereby lacking conservation, or TFBSs in TEs since these regions are usually masked when determining sequence conservation (Haudry *et al*., 2013; Siepel *et al*., 2005; Van De Velde *et al*., 2014). Among the included CNS methods, we show that the CNS method by (Song *et al*., 2021) provides the best results. By covering a smaller evolutionary distance than other methods (only species within the *Andropogoneae* tribe), it might be able to detect more recent TFBSs, enhancing recall, while maintaining a good precision. Following the CNS methods, we find chromatin accessibility to be a better approach to identify functional CREs. However, ACRs might miss the TFBSs of pioneering TFs, which don’t need accessibility to perform their function (Lai *et al*., 2021; Strader *et al*., 2022).

Organ- or tissue-specificity in our datasets should be taken into account when interpreting our results. Our benchmark puts forward UMRs as the most informative for predicting TFBSs. However, a reason for the superior performance of UMRs might be that they were profiled in maize leaf, similar to our benchmark’s ChIP-seq gold standard. While UMRs are considered relatively stable across organs/tissues, some degree of specificity remains, leading to a potential overestimation of their performance (Crisp *et al*., 2020). The ACRs we compiled combine accessibility data from different organs and tissues, and are therefore not expected to be strongly specific. Notably, a single-cell ATAC-seq dataset was included (Marand *et al*., 2021), representing a high-resolution set of ACRs of different organs and cell types. However, as not every cell type, developmental stage, and condition is included in our ACR compendium, this dataset is expected to still exhibit some context-specific biases (e.g., missing of CREs associated with specific *in vivo* contexts). In contrast, genome conservation is fully context-independent. The leaf-specificity of the gold standard ChIP-seq experiments could result in an underestimation of CNS performance when considering all CREs in the maize genome. Indeed, we do find some evidence for this hypothesis in the additional benchmark against the less organ-specific UMR silver standard, where CNSs are outperforming ACRs. This indicates that CNSs might not necessarily be less informative than ACRs when evaluated in a more context-independent manner.

Assembling the iCREs raised the opportunity to extend MINI-AC’s functionality to infer context-specific GRNs using the iCREs and a list of genes. Currently, the state-of-the-art methods to infer GRNs have a strong focus on exploiting single-cell (multi-)omics datasets because they offer an unprecedented level of resolution (Bravo González-Blas *et al*., 2023; Ferrari *et al*., 2022; Fleck *et al*., 2023; Jiang *et al*., 2022; Kamimoto *et al*., 2023). However, those protocols are still not optimized for every model species and/or experimental condition, and they are also expensive, which might make them suboptimal or unnecessary, depending on the biological question being addressed. Conversely, the inference of GRNs from a specific list of genes can aid in highlighting candidate regulators in a simple and straightforward manner. Such tools exist, like TF2Network, TDTHub, ConnecTF, or PlantRegMap (Brooks *et al*., 2021; Grau and Franco-Zorrilla, 2022; Kulkarni *et al*., 2018; Tian *et al*., 2020). However, they test motif enrichment by applying a hypergeometric or Fisher’s exact test to TFBSs in pre-defined genomic windows adjacent to genes (e.g., upstream promoter region), which misses distal CREs. In our iCREs-based framework the motif enrichment encompasses the whole non-coding genome, but reduces potential false positive TFBSs. We showed that this increases the performance in the prediction of expected regulators. We found that the “maxF1” iCREs set, which is the overlap between the UMRs and the ACRs, has the best agreement (F1 metric) with the leaf ChIP-seq gold standard. The “all iCREs” set, containing all UMRs, ACRs and CNSs, has a poorer agreement with the gold standard. However, when used to infer GRNs, it marginally improves predictions of enriched binding sites for the differentially expressed regulators in an input set of genes. This, however, is not completely unexpected, given that the gold standard is derived from mesophyll cells in normal conditions, and we predict drought GRNs for multiple organs. Thus, the aforementioned elements unique to the “all” iCREs set might contain additional relevant CREs not captured by the ChIP-seq mesophyll gold standard. Integration of a leaf eQTL dataset showed the “all iCREs” have a high enrichment for lead SNPs of drought-specific eQTLs, confirming the power of the iCREs to demarcate regulatory DNA. Furthermore, the drought-responsive GRNs, constructed using drought DE genes and the “all iCREs”, significantly overlapped with the eQTL regulatory interactions, enhancing the identification of novel regulators and target genes conferring organ-specific transcriptional regulation.

The regulatory role of PIF/Harbinger TEs found in our analysis has been previously explored and confirmed in *Brassica oleracea*. In this species, insertions of TEs belonging to this superfamily altered the expression of a TF responsible for purple traits in cultivars. Additionally, reporter assays showed that promoter sequences containing this TE insertion induced higher expression than those that do not (Li *et al*., 2024). These results are in line with our findings, where we observed that genes close to both TEs within and outside iCREs have higher expression levels than those close to only TEs outside iCREs. This indicates that those TEs have a promoting effect on gene expression rather than a repressing effect, which is in agreement with a study revealing an important role for PIF/Harbinger-driven enhancers in the regulation of husk development (Fagny *et al*., 2020). The role of TEs on stress gene regulatory networks in *Arabidopsis thaliana* and tomato has previously been assessed, where enriched motifs were found in different TE superfamilies, similar as in our study; e.g., WRKY TFs in Mariner TEs (Deneweth *et al*., 2022). In addition, the contribution of TEs to the stress response in maize has been studied before (Le *et al*., 2014; Makarevitch *et al*., 2015; Roquis *et al*., 2021), as well as the potential regulatory role of maize TEs within ACRs (Noshay *et al*., 2021; Zhao *et al*., 2018). Our findings provide support for previously proposed models where TEs play a role in rewiring GRNs (Britten and Davidson, 1971; Feschotte, 2008; Rebollo *et al*., 2012). While this phenomenon has been observed in animals and some plants (Batista *et al*., 2019; Kunarso *et al*., 2010; Lynch *et al*., 2011; Wang *et al*., 2007), our results hypothesize that TIR TEs play a similar role in wiring maize stress response networks.

In our study, the assumption that one iCRE controls its closest gene fails to capture long-range regulatory interactions, where adjacent genes are skipped, or cases where one iCRE controls more than one gene. Eventually, the generation of high-quality Hi-C (Lieberman-Aiden *et al*., 2009) and ChIA-PET (Fullwood *et al*., 2009) datasets in maize for different conditions would allow integrating them in our framework to resolve these limitations. Nonetheless, in maize, these techniques have only been applied in few conditions, limiting their general usage for predictions in other organs/tissues, conditions, or genotypes, given that these long-range interactions can be tissue-specific (Marand *et al*., 2021). Moreover, the innovation of a gold standard of CREs in maize that is context-independent and contains minimum false positives could improve the data-driven optimization and integration of iCREs. As demonstrated using drought eQTL data, iCREs hold the potential to be integrated with data of natural genetic variation associated with phenotypic changes, to explore the role of non-coding variants within TFBSs and GRNs in determining complex traits.

In summary, our study highlights the potential of integrating data derived from different methods to aid regulatory genome analysis in different biological contexts. By combining epigenetics and genomics data, we generated a comprehensive collection of maize CREs, supporting the scientific community working in maize, and highlighting the complexity and intricacies of regulatory DNA in complex plant genomes.

## Materials and Methods

### CNS inference

The genomic coordinates of the CNSs from (Song *et al*. 2021) were downloaded from https://genome.cshlp.org/content/suppl/2021/06/22/gr.266528.120.DC1. The pairwise CNS coordinates files were concatenated and merged (*bedtools merge;* version 2.2.28) (Quinlan and Hall, 2010). Coordinates were then converted from version 4 to version 5 of the maize B73 reference genome using the UCSC liftOver tool (Hinrichs *et al*., 2006) (default parameters) and a version 4 to version 5 chain file available via maizeGDB (www.maizegdb.org) (Woodhouse *et al*., 2021). Genome-wide maize CNSs predicted by funTFBS were downloaded via the Plantregmap portal (http://plantregmap.gao-lab.org) (Tian *et al*., 2020) and converted from version 3 to version 4 using the online Assembly Converter tool of Ensembl plants for maize (Martin *et al*., 2023) with default parameters. Subsequently, they were converted from version 4 to version 5 using liftOver as described above. CNSs predicted by the Conservatory project (Hendelman *et al*., 2021) were downloaded from https://conservatorycns.com/dist/pages/conservatory/index.php as maize B73 version 5 coordinates.

BLSSpeller (De Witte *et al*., 2015) and *msa_pipeline* (Wu *et al*., 2022) were run using a species set covering the PACMAD clade, including species *Cenchrus purpureus*, *Oropetium thomaeum*, *Saccharum spontaneum*, *Setaria italica, Sorghum bicolor*, *Zea mays* and *Zoysia japonica* ssp. *Nagarizaki*. To obtain orthologous groups and a species tree, OrthoFinder (Emms and Kelly, 2019) version 2.5.4 was used with default settings, using proteome files consisting of longest transcripts as input (retrieved from PLAZA version 5.0) (Van Bel *et al*., 2022). The resulting orthologous groups were filtered for groups that contain genes of at least two species, including maize B73. The genome FASTA files used by BLSSpeller and msa_pipeline were retrieved from PLAZA version 5.0. Repeats were masked using a k-mer-based approach as described by (Song *et al*., 2021), using KAT (Mapleson *et al*., 2017). For BLSSpeller, a search space of 2 kb upstream of the translation start site, 1 kb downstream of the translation end site of all the genes, including the introns was used. Next, the coding sequences were masked. BLSSpeller (source code available at https://bitbucket.org/dries_decap/bls-speller-spark) assembly version 1.1 was used in alignment-free mode on the upstream, downstream and intronic regions separately, using 8 bps as motif length, using the full IUPAC alphabet of degenerate nucleotides, and a maximum of 3 degenerate characters in a motif. A confidence score cutoff of 85 was used, the minimal number of conserved orthologous group counts was set to 25, and the used range of BLS thresholds was [50, 60, 70, 75, 80, 85, 90, 92, 95, 97, 99]. *Msa_pipeline* was used as described by (Wu *et al*., 2022) (available at https://bitbucket.org/bucklerlab/msa_pipeline), using their relaxed LAST parameter set. The initial LAST step using lastdb was run with parameter -u RY32 to decrease runtime. After pairwise alignment, alignments were chained and netted (Kent *et al*., 2003) followed by multiple genome alignment by ROAST (https://www.bx.psu.edu/~cathy/toast-roast.tmp/README.toast-roast.html). *Zea mays* was used as the reference species. If a comparator species was polyploid (*Triticum aestivum*, *Triticum turgidum* and *Saccharum spontaneum*), each subgenome was considered as a separate species. An adaptation of the *msa_pipeline*’s conservation calling procedure was used, where conserved elements were called per chromosome using GERP++ (Davydov *et al*., 2010) with default parameters. Conservation scores were generated using *gerpcol*, and conserved elements using *gerpelem*.

### Experimental regulatory datasets

The dataset containing ChIP-seq summits of 104 TFs in maize was obtained from (Tu *et al*., 2020), and a region of 10 bp upstream and 10 bp downstream around a summit (total of 21 bp) was considered the peak region. The peak regions of all individual TFs were concatenated and merged (*bedtools merge*). The UMRs genomic coordinates were downloaded from the supplementary dataset 3 of (Crisp *et al*., 2020). Publicly available ACR datasets were collected from GEO and PlantCADB (https://bioinfor.nefu.edu.cn/PlantCADB/; (Ding *et al*., 2022)). Thirteen high-quality ACR datasets were selected by omitting negative control experiments (naked DNA) and removing datasets that, similar to negative control samples, showed a high fraction (>70%; Figure S1) of distal regions (further than 2 kb from a gene), indicating low quality. Processed BED files were collected from GEO for GSE120304 (leaf and ear) (Ricci *et al*., 2019) and GSE155178 (the provided peak file was converted to BED) (Marand *et al*., 2021). The datasets GSE128434 (Lu *et al*., 2019), GSE85203 (Lu *et al*., 2017), GSE94291 (Oka *et al*., 2017), GSE97369 (Burgess *et al*., 2019), PRJNA382414 (Zhao *et al*., 2018), PRJNA391551 (Dong *et al*., 2017), PRJNA518749 (Tu *et al*., 2020), and PRJNA599454 (Sun *et al*., 2020) were downloaded from PlantCADB (Ding *et al*., 2022). Replicates per experiment were combined by intersecting them with *bedtools intersect*. All the ACR datasets were concatenated and merged (*bedtools merge*) into a single cross-tissue ACR dataset. The coordinates of all the datasets were converted from version 4 to version 5 of the maize B73 reference genome using the UCSC liftOver tool (default parameters) and a version 4 to version 5 chain file available via maizeGDB (www.maizegdb.org).

### Performance evaluation

The comparison of a query and a reference dataset containing genomic coordinates was done by computing, precision, recall and F1 as follows:

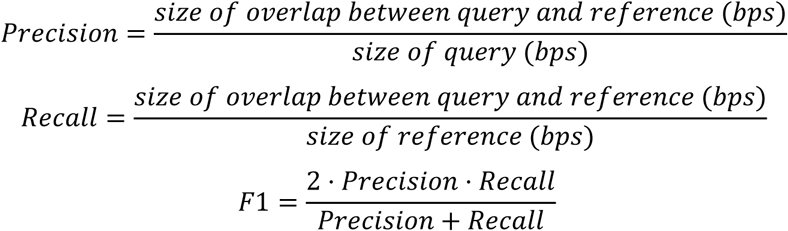

Performance was calculated, considering only the non-coding genome by first intersecting (*bedtools intersec*t) query and reference regions with the non-coding genome (GFF downloaded from PLAZA Monocots 5.0), after which the formulas above were applied.

### Computation of enrichment between sets of genomic regions

The enrichment of a query set of genomics regions (for example, the CNSs) in a target set of genomic regions (for example, the ChIP-seq peaks) was tested by shuffling the latter 1,000 times across the maize non-coding or full genome. Then, the observed overlap (real overlap) between the genomic sets was compared to the overlap of the shuffled sets (overlap expected by chance). A p-value was then computed as the number of times the overlap with any of the 1,000 shuffled sets was higher than the real overlap. Enrichment fold was computed as the real overlap divided by the median of the overlap expected by chance (median of the 1,000 shuffling events).

### Construction of iCRE sets

The “max F1 iCREs” set was obtained by intersecting (*bedtools intersect*) the UMRs with the cross-tissue ACR dataset (see above). The “all iCREs” set was obtained by concatenating and merging (*bedtools merge*) all UMRs, ACRs and CNSs of (Song *et al*. 2021). Last, the coding sequences as defined in the maize version 5 annotation and a small (59 kb) set of high confidence predicted t-RNAs (available via http://gtrnadb.ucsc.edu/), were subtracted (*bedtools subtract*) from both iCRE sets. The genomic annotation of the regions was done with the maize GFF downloaded from PLAZA Monocots 5.0.

### Processing of the gene expression atlas of maize in drought conditions

A manual curation of *Zea mays* drought RNA-seq datasets was done using CurSE (Vaneechoutte and Vandepoele, 2019) with default parameters. In total, 20 studies, summarized in Table S2, were selected. A total of 434 samples and their corresponding FASTQ files were downloaded using *fastq-dump -I –split-files -gzip* (version 3.0.0 https://github.com/ncbi/sra-tools) and processed using the nf-core/rnaseq (Ewels *et al*., 2020) pipeline to process RNA-seq data. The input sample sheets were obtained using the *fastq_dir_to_samplesheet.py* script provided in the nf-core/rnaseq repository. The transcriptome, genome FASTA, and GTF files corresponding to the maize genome version 5 were downloaded from PLAZA Monocots 5.0 (Van Bel *et al*., 2022). Only the stages 1 (pre-processing) and 3 (pseudo-alignment and quantification) of the nf-core/rnaseq pipeline were executed using the default parameters, except for the specification of Salmon as pseudo-aligner for quantification. The index file for Salmon was obtained using *salmon index* with default parameters and without decoys (Patro *et al*., 2017). Samples with a mapping rate of less than 65% were discarded, as well as outlier samples within each study with a mapping rate of less than 70% (Figure S4A). Outliers in each study were samples with a mapping rate below this value: Q1 - 1.5 × (Q3-Q1) (Q1 and Q3 being the first and third quartile, respectively). The contrasting control or treatment samples of the ones discarded due to low mapping rate were also removed, leaving a total of 375 samples (141 after processing replicates).

### Selection of differentially expressed genes in response to drought

Differential expression analysis was performed using *edgeR* (Robinson *et al*., 2010) with default parameters. Genes with less than one transcript per million (TPM) in at least two samples were discarded. Next, DEGs were inferred using *glmQLFit* and *glmQLFTest*, with the multiple testing correction and the false discovery rate (FDR) calculations being done with the Benjamini-Hochberg procedure (Benjamini and Hochberg, 1995). For each contrast, the genes with an FDR < 0.05 and an absolute log-fold change > 1 were considered differentially expressed. Of the 80 contrasts in this analysis, a large proportion did not yield any DEG, or very few (Figure S4B). Contrasts yielding less than 100 up- or down-regulated genes were considered low quality and therefore discarded, leaving a total of 55 contrasts. The activation and repression consistency scores of the DEG sets were computed as the number of contrasts in which they are up- or down-regulated, divided by the total number of contrasts in which they are differentially expressed.

### iCREs-based gene regulatory networks inference

The “maxF1” and “all” iCREs sets were individually annotated to the closest gene using *bedtools closest -a $icres -b $genes -t all*. MINI-AC was employed to perform motif enrichment and GRN inference (Manosalva Pérez *et al*., 2024) on the iCREs associated to each drought-responsive gene set. MINI-AC was run with the genome-wide mode on the maize genome version 5, and with the “absolute base pair count” option set to “true”. An iCREs-based MINI-AC feature was added to the GitHub repository (https://github.com/VIB-PSB/MINI-AC/tree/main) to infer context-specific GRNs in maize given a list of genes. The coordinates of the neighboring non-coding regions and introns of each gene were obtained from the maize GFF file from PLAZA 5.0 and were then overlapped with the “medium” promoter file provided in the MINI-AC GitHub repository (https://github.com/VIB-PSB/MINI-AC/blob/main/data/zma_v5/zma_v5_promoter_5kbup_1kbdown_sorted.bed). This file was used as input for MINI-AC, using the same parameters as specified above, to perform promoter-based motif enrichment. To compare the promoter-based and the iCREs-based motif enrichment approaches, the precision was computed as the number of enriched motifs associated with differentially expressed TFs divided by the total number of enriched motifs. The recall was computed as the number of enriched motifs associated with differentially expressed TFs divided by the total number of motifs associated with differentially expressed TFs. F1 was computed as the harmonic mean of precision and recall. The AUPRC was computed using the function *metrics.auc* (default parameters) from the Python library *sklearn* (Pedregosa *et al*., 2011).

### eQTL analysis

The eQTL data was obtained from (Liu *et al*., 2020), Table S2. Drought-specific eQTLs were extracted as eQTLs that were found in either of the two drought experiments, but not in the well-watered condition. Coordinates were converted from maize B73 version 4 to version 5 using the UCSC liftOver tool (Hinrichs *et al*., 2006) (default parameters), using a chain file available via maizeGDB (www.maizegdb.org). Enrichment of the eQTL lead SNPs was performed as described earlier in the section describing enrichment between sets of genomic regions, here shuffling the iCREs 1,000 times within the non-coding genome to create the background model. Significance in overlap between a MINI-AC GRN and the eQTL GRN was determined using a background set of 1,000 randomized GRNs, constructed by randomly drawing a TF and a target gene from the MINI-AC GRN, creating random edges. A p-value for the overlap was computed as the number of times the overlap with a shuffled GRN was higher than the real overlap.

### Transposable elements analysis and chromatin states enrichment and STARR-Seq

The coordinates of the maize transposable and repetitive elements in the genome version 5 were downloaded from MaizeGDB (https://download.maizegdb.org/Zm-B73-REFERENCE-NAM-5.0/Zm-B73-REFERENCE-NAM-5.0.TE.gff3.gz) (Woodhouse *et al*., 2021). The coordinates were split into TE superfamilies based on the third column value (feature) of the GFF file. The genomic coordinates of the chromatin states were downloaded from the plant chromatin state database (http://systemsbiology.cau.edu.cn/chromstates/download.php; (Liu *et al*., 2018)) and converted from the AGPv3 genome version to the AGPv4 genome version, and then to the B73 RefGen v5 genome version using liftOver with default parameters (Hinrichs *et al*., 2006). The chain file used by liftOver was downloaded from Ensembl (Martin *et al*., 2023) (https://ftp.ensemblgenomes.ebi.ac.uk/pub/plants/release-55/assembly_chain/zea_mays/AGPv3_to_B73_RefGen_v4.chain.gz and https://ftp.ensemblgenomes.ebi.ac.uk/pub/plants/release-55/assembly_chain/zea_mays/B73_RefGen_v4_to_Zm-B73-REFERENCE-NAM-5.0.chain.gz). The TEs fully contained within iCREs (*bedtools intersect -a $tes -b $icres - f* 1) were split based on the TE superfamily. The enrichment of TEs, TEs superfamilies, and other genomic region sets in the iCREs and the chromatin states was done as described previously in this section. The shuffling was done across the non-coding genome for the iCREs enrichment and across the whole genome for the chromatin states enrichment. For visualization purposes, the enrichment metrics were integrated into the π-value, which is the −log10(p-value) × enrichment fold. The p-value was adjusted for multiple testing using the Benjamini–Hochberg method with significance level of 0.01. The processed STARR-seq data was obtained from GEO (GSE120304) (Ricci *et al*., 2019), and the raw input library reads were downloaded from SRA (SRR10964903). Raw reads were trimmed with Trimmomatic v0.36 (SLIDINGWINDOW:3:20 LEADING:0 TRAILING:0 MINLEN:30) (Bolger *et al*., 2014) and aligned to the B73 RefGen v5 genome (PLAZA Monocots 5.0 (Van Bel *et al*., 2022)) using Bowtie v0.12.7 (*bowtie -t -v 1 -X 2000 --best --strata -m 1 -S*) (Langmead *et al*., 2009). Mapped input reads were converted to a BED file (*bamtobed*) and then merged (*merge*) using bedtools v2.2.28 with default parameters. Within these input regions, TEs got assigned an enhancer activity score by averaging the enhancer activity of each base pair in the TE. The increase in enhancer activity for TEs within iCREs, compared to outside iCREs, was tested using a one-sided Mann-Whitney U test, corrected for multiple testing using the Benjamini–Hochberg method.

To perform functional analysis, TEs were annotated to their closest gene (*bedtools closest -a $tes_in_icres -b $genes -t all -d*), and GO enrichment was performed using a gene-GO annotation file of the maize genome version 5 (https://github.com/VIB-PSB/MINI-AC/blob/main/data/zma_v5/zma_v5_go_gene_file.txt). The enrichment was calculated using a hypergeometric test and the p-value was corrected using the Benjamini-Hochberg procedure.

The iCREs linked to genes that were associated with PIF-Harbinger and Tc1-Mariner superfamilies, and that yielded enrichment in the GO terms “defense response to bacterium”, “regulation of flower development”, and “response to salt stress” were used for motif enrichment using MINI-AC with the same parameters as described earlier in this section. To evaluate the expression levels of genes associated with TEs within iCREs, the TPMs per gene and per sample of the drought gene expression atlas mentioned earlier in this section were employed. The distance to each gene was obtained as previously mentioned using *bedtools closest*.

## Author contributions

KV, NMP, and JS conceived and designed the research. JS performed the benchmark and complementarity analysis of CRE profiling methods. NMP and JS performed the analysis of the gene regulatory networks and transposable elements. ID ran *msa_pipeline*. AMF processed the gene expression atlas of maize drought. NMP, JS, and KV wrote the manuscript with input from the rest of the authors.

## Declaration of generative AI and AI-assisted technologies in the writing process

During the preparation of this work the authors used ChatGPT in order to rephrase content written by the authors. After using this tool/service, the authors reviewed and edited the content as needed and take full responsibility for the content of the publication.

## Acknowledgements

We would like to thank Dries Decap and Jan Fostier for their support and help running BLSSpeller. We also thank Chrystian Camilo Sosa Arango for sharing an RNA-seq differential expression pipeline, used to call drought responsive DEGs. This work was supported by a Bijzonder Onderzoeksfonds grant from Ghent University (grant agreement: BOF24Y2019001901) to NMP and by The Research Foundation - Flanders (FWO; Odysseus II G0D0515N) to JS.

## Supplementary datasets

Files containing the genomic coordinates of the conserved non-coding sequences (BLSSpeller, msa_pipeline, funTFBS, Song-2021, and Conservatory), the accessible chromatin region compendium (ACRs) and the unmethylated regions (UMRs) employed to generate the integrated cis-regulatory-elements (iCREs), and both the “all iCRE” and “maxF1 iCRE” sets, are available at https://doi.org/10.5281/zenodo.15143951. These datasets are available both in version 4 and 5 of the maize genome.

---

**Figure S1**

**Figure S1. Length distribution, genome coverage, and genomic feature distribution of the collected datasets with putative cis-regulatory elements (CREs), and transposable and repetitive elements.** (A) The length distribution of the genomic regions in a dataset (indicated on the x-axis at the bottom), annotated with the number of regions (n) in the dataset. (B) The coverage of genomic features by the different regions of each dataset, i.e., what percentage of a given genomic feature is covered by the dataset. Datasets are annotated with their total size in megabases (Mb). Genomic features are annotated with their total size in Mb and as a percentage of the total genome size. (C) The genomic feature distribution in which the dataset regions are located, i.e., what percentage of a dataset is located in each genomic feature.

**Figure S2**

**Figure S2. Precision-recall plot of individual CNS detection and experimental CRE-profiling methods, using a silver standard.** Precision-recall plot where performance is calculated against a cross-tissue ACR dataset (A), or a UMR dataset (B).

**Figure S3**

**Figure S3. Benchmark and comparison of the conserved non-coding sequences (CNS) detection methods.** An UpSet plot showing the sizes of the full datasets on the left and the sizes of the unique subsets on the top. Precision, recall and F1 are calculated using a ChIP-seq gold standard for each of the individual subsets in a heatmap below the size bars. The UpSet plot is sorted on the precision of the subsets, so that the subset with the most correct predictions is shown on the left. Precision, recall and F1 are also calculated for different ensemble sets, created by starting from the subset with the highest precision (most left) and progressively adding individual subsets with the next best precision, until all regions are added together (most right).

**Figure S4**

**Figure S4. Quality control metrics of the maize drought gene expression atlas processing.** (A) Box plot showing the distribution of the mapping rates of the samples per study. (B) Box plot showing the distribution of number of differentially expressed genes in the samples, per study.

**Figure S5**

**Figure S5. Distribution of the activation and repression consistency scores of the tissue-specific “marker” drought-responsive genes.**

**Figure S6**

**Figure S6. Complete UpSet plot showing the number of common and shared drought-responsive genes per tissue.** All intersection sets are shown, and they are sorted by size.

**Figure S7**

**Figure S7. Complete clustered heatmap showing the motif enrichment rank of differentially expressed transcription factors (TFs) in the inferred gene regulatory networks (GRNs).** TFs (y-axis) are shown for the “all iCREs”-based GRNs, inferred using sets of iCREs that are associated with tissue-specific (x-axis) up-regulated genes in response to drought. The row annotations correspond to the activation and repression consistency score bins and the TF family.

**Figure S8**

**Figure S8. Clustered heatmap of gene ontology (GO) enrichment for target genes of predicted regulators in the inferred drought-responsive gene regulatory networks.** The −log10(q-value) of GO (x-axis) enrichments are displayed among the predicted target genes of up- and down-regulated transcription factors (TFs) in response to drought in leaf (y-axis). The row annotations indicate the activation and repression consistency score of the TFs, and the TF family. The column annotation corresponds to the gene set (up- or down-regulated marker genes in response to drought in leaf) that yielded the indicated GO enrichment, and the number of target genes in that GO category. For visualization purposes, the enriched GO terms with 5 or fewer target genes were discarded, as well as TFs with 3 or less GO terms enriched.

**Figure S9**

**Figure S9. Chromatin states enrichment per TE superfamily and repetitive elements type.** Clustered heatmap showing the chromatin state (y-axis) enrichment π-value of different TE subsets split per TE superfamily or repetitive elements type (x-axis). The columns are annotated based on the TE superfamily/repetitive elements type, and based on the TE subset (all TEs, TEs within iCREs, and TEs outside iCREs).

**Figure S10**

**Figure S10.** Distribution of the distance to the closest gene for the indicated sets of genomic regions.

## Tables

**Table S1**

**Table S1. Performance metrics and genome coverage of individual datasets.** Performance is calculated using a ChIP-seq gold standard. The adjusted p-value represents the significance of overlap between the individual dataset and the gold standard.

**Table S2**

**Table S2. List of the maize drought studies employed in this publication, their metadata, and characteristics.**

**Table S3**

**Table S3. Summary statistics of all gene regulatory networks constructed in this study.**

**Table S4**

**Table S4. Performance of the “promoter-based”, “all iCREs-based”, and “max F1 iCREs” gene regulatory networks.** The two performance metrics (F1 and area under the precision recall curve) measure how well each method predicts differentially expressed regulators, given a set of genes.

**Table S5**

**Table S5. Gene ontology (GO) enrichment of the genes associated with different repetitive element types and transposable element (TE) superfamilies that are found within iCREs.**

**Table S6**

**Table S6. Number of, median value of the transcripts per million, and p-value of the genes associated with transposable elements (TEs) within, outside, and within and outside iCREs.** The computations were binned per distance to the closest gene.

**Table S7**

**Table S7. STARR-seq enhancer activity of transposable element (TE) superfamilies within and outside iCREs.**

**Table S8**

**Table S8. Enrichment results of ChIP-seq peaks in PIF Harbinger and Tc1-Mariner TEs in iCREs and outside iCREs.**

## Datasets

**Dataset S1**

**Dataset S1. Genomic coordinates of the conserved non-coding sequence datasets, accessible chromatin region compendium, unmethylated regions, and iCREs.** Both the “all iCREs” and “max F1 iCREs” are included. All datasets are available both in version 4 and 5 of the maize genome.

**Dataset S2**

**Dataset S2. Edge list files of all gene regulatory networks constructed for this study.**

**Dataset S3**

**Dataset S3. Gene ontology enrichment output of “all iCRE”-based gene regulatory networks.**

