## Supplemental Figures for "A map of integrated cis-regulatory elements enhances gene regulatory analysis in maize"

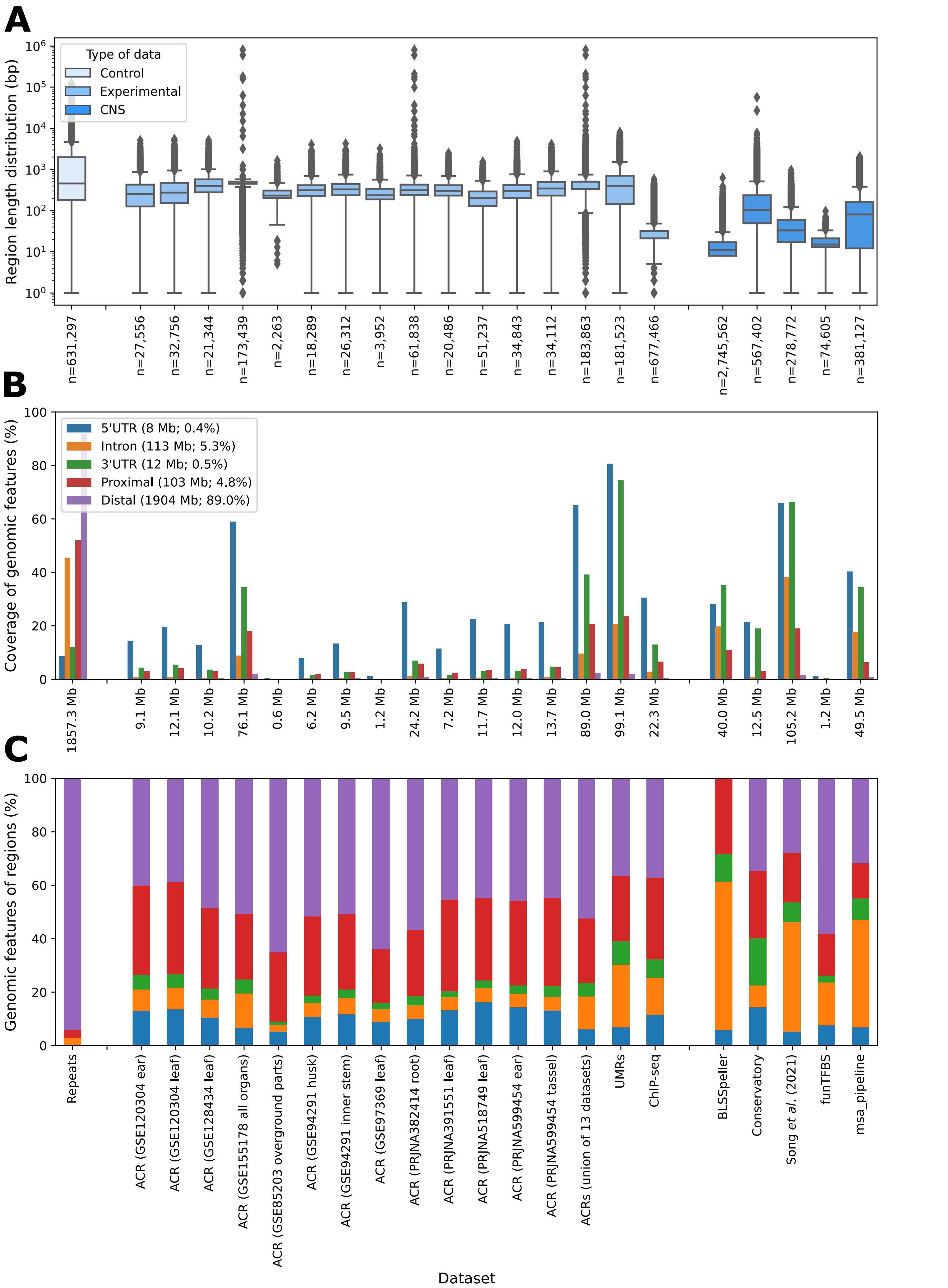


**Supplementary Figure 1. Length distribution, genome coverage, and genomic feature distribution of the collected datasets with putative cis-regulatory elements (CREs), and transposable and repetitive elements.** (A) The length distribution of the genomic regions in a dataset (indicated on the x-axis at the bottom), annotated with the number of regions (n) in the dataset. (B) The coverage of genomic features by the different regions of each dataset, i.e., what percentage of a given genomic feature is covered by the dataset. Datasets are annotated with their total size in megabases (Mb). Genomic features are annotated with their total size in Mb and as a percentage of the total genome size. (C) The genomic feature distribution in which the dataset regions are located, i.e., what percentage of a dataset is located in each genomic feature.

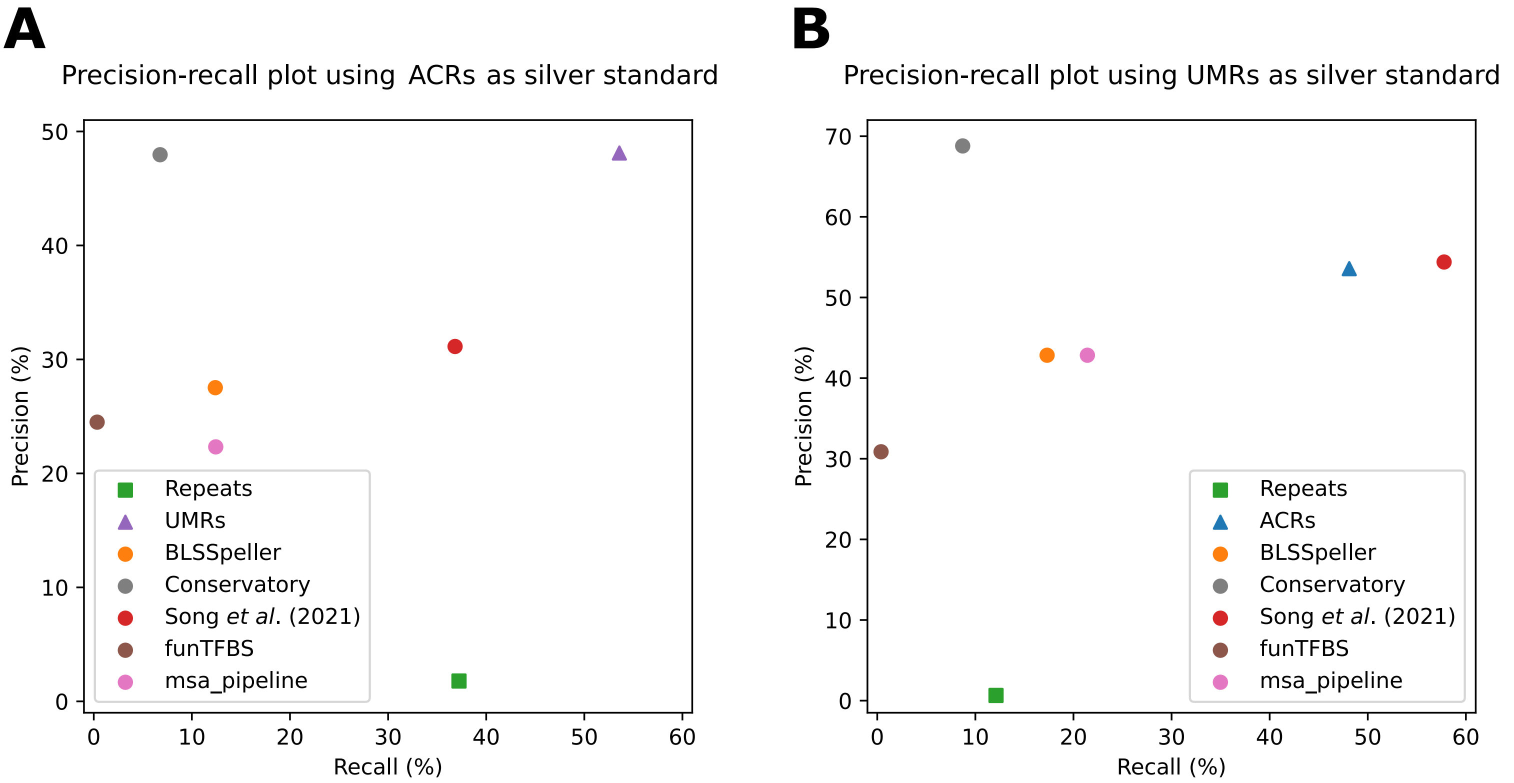


**Supplementary Figure 2. Precision-recall plot of individual CNS detection and experimental CRE-profiling methods, using a silver standard.** Precision-recall plot where performance is calculated against a cross-tissue ACR dataset (A), or a UMR dataset (B).

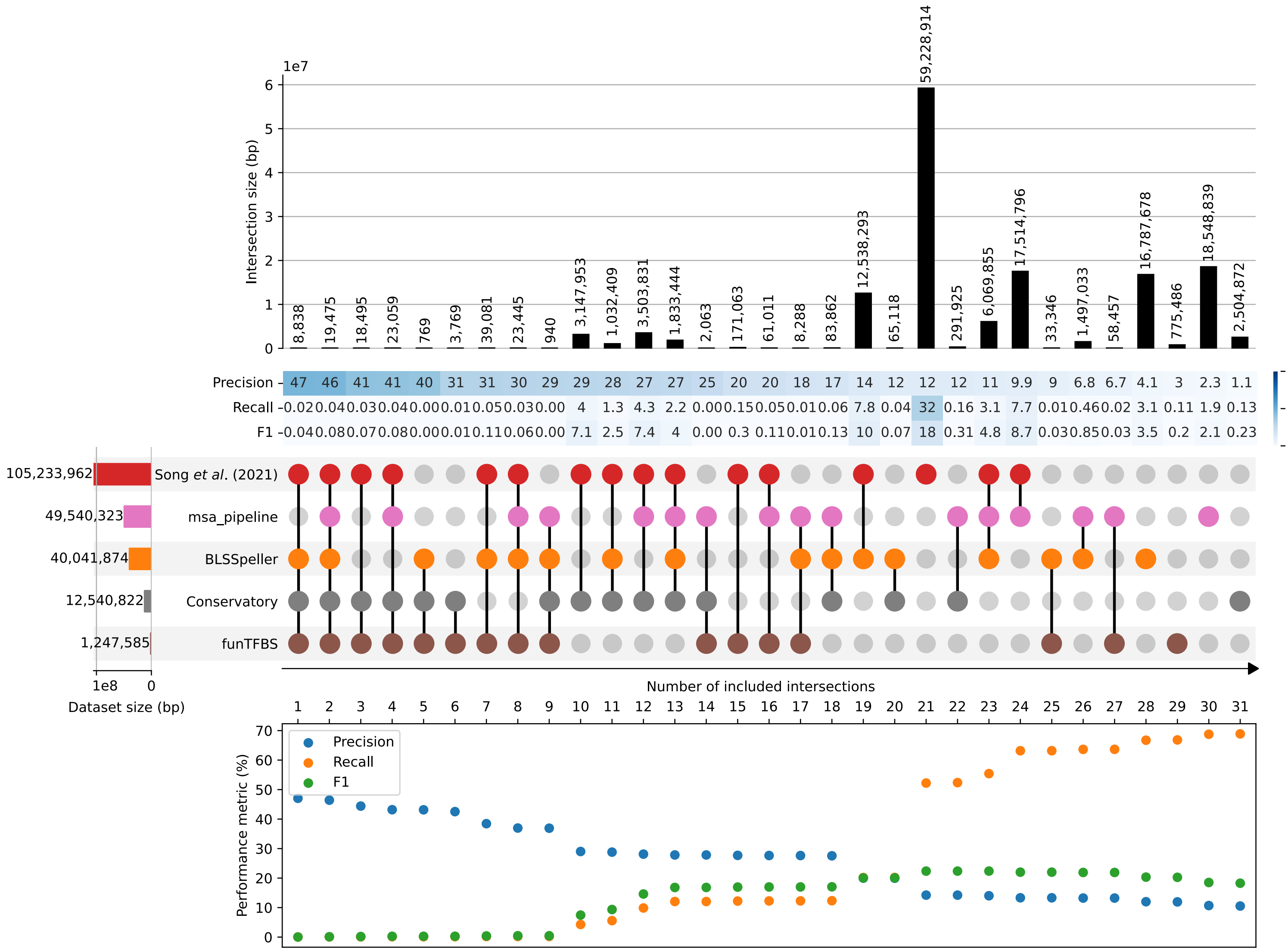


**Supplementary Figure 3. Benchmark and comparison of the conserved non-coding sequences (CNS) detection methods.** An UpSet plot showing the sizes of the full datasets on the left and the sizes of the unique subsets on the top. Precision, recall and F1 are calculated using a ChIP-seq gold standard for each of the individual subsets in a heatmap below the size bars. The UpSet plot is sorted on the precision of the subsets, so that the subset with the most correct predictions is shown on the left. Precision, recall and F1 are also calculated for different ensemble sets, created by starting from the subset with the highest precision (most left) and progressively adding individual subsets with the next best precision, until all regions are added together (most right).

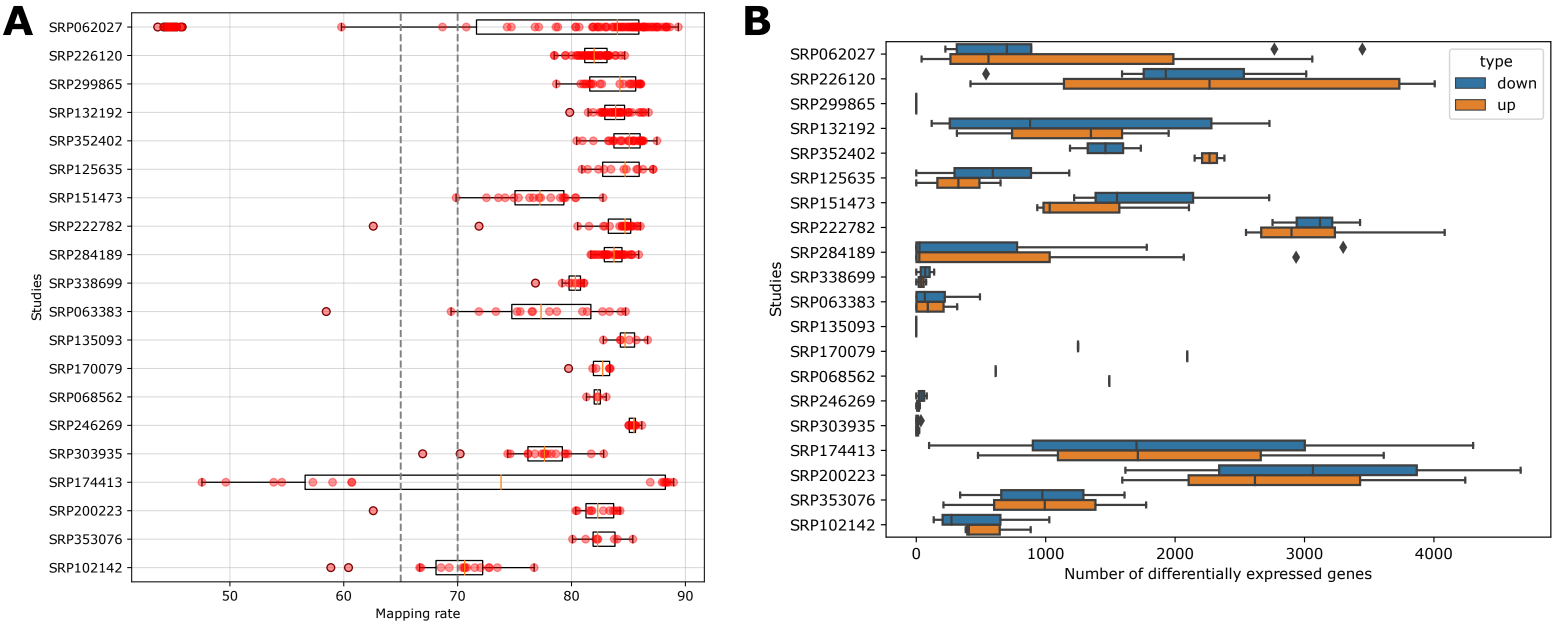


**Supplementary Figure 4. Quality control metrics of the maize drought gene expression atlas processing.** (A) Box plot showing the distribution of the mapping rates of the samples per study. (B) Box plot showing the distribution of number of differentially expressed genes in the samples, per study.

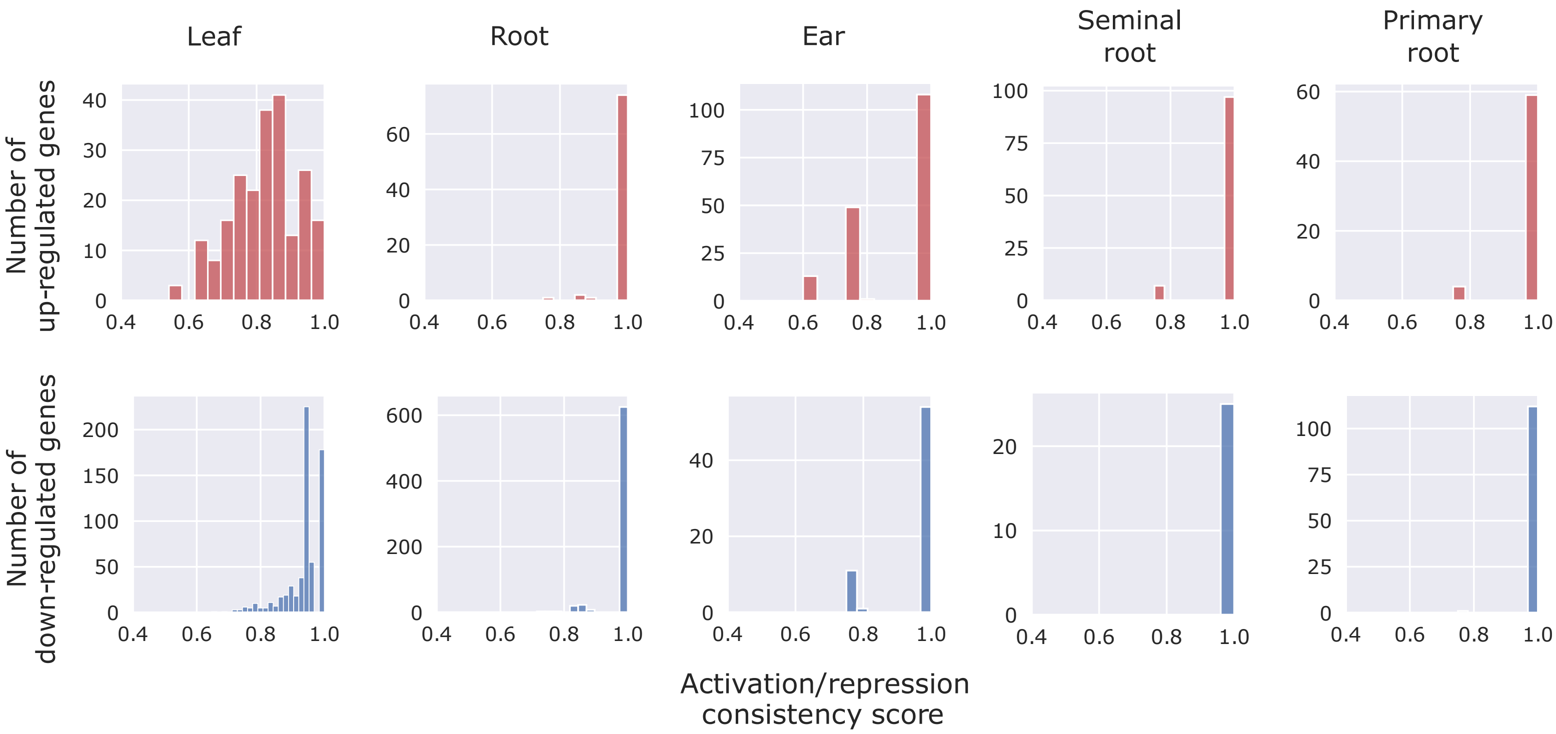


**Supplementary Figure 5. Distribution of the activation and repression consistency scores of the tissue-specific "marker" drought-responsive genes.**

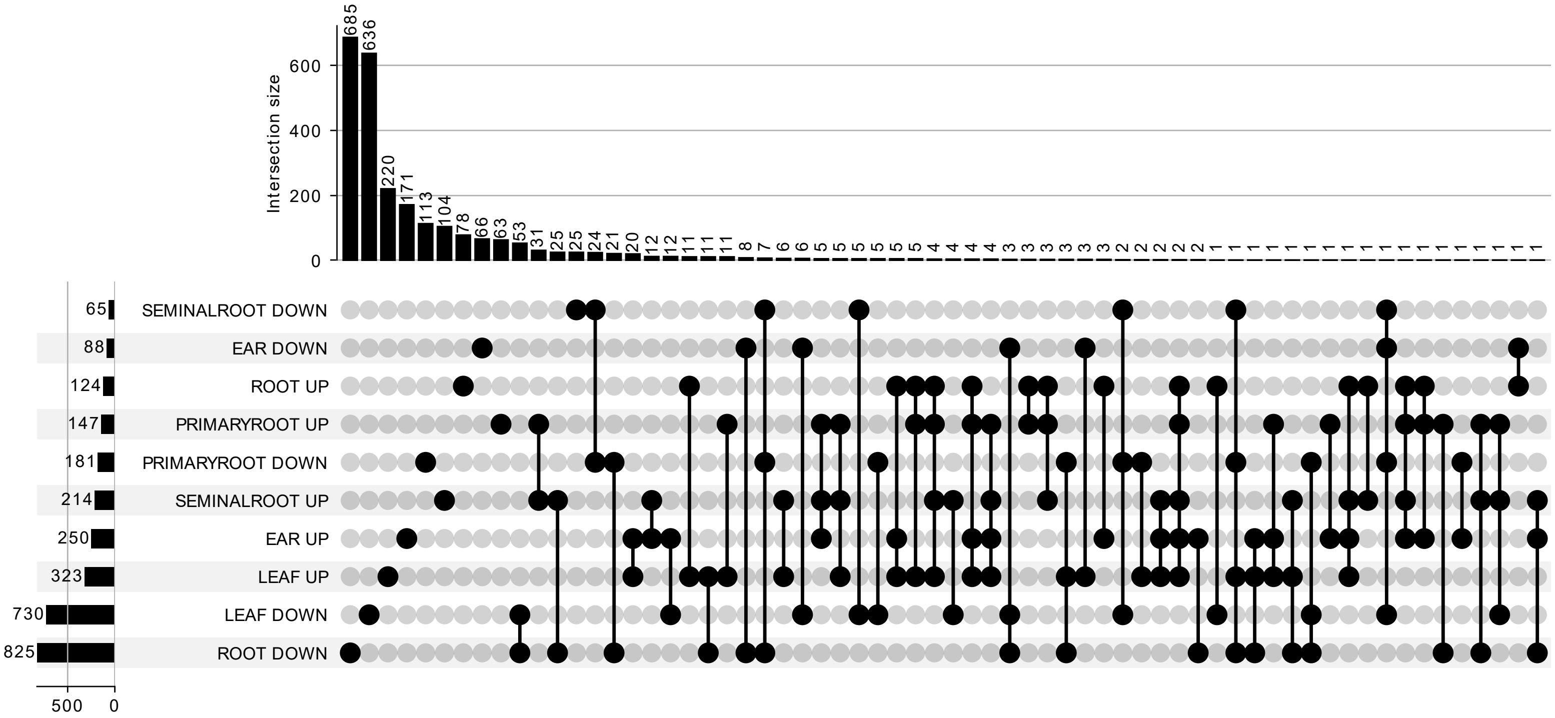


**Supplementary Figure 6. Complete UpSet plot showing the number of common and shared drought-responsive genes per tissue.** All intersection sets are shown, and they are sorted by size.

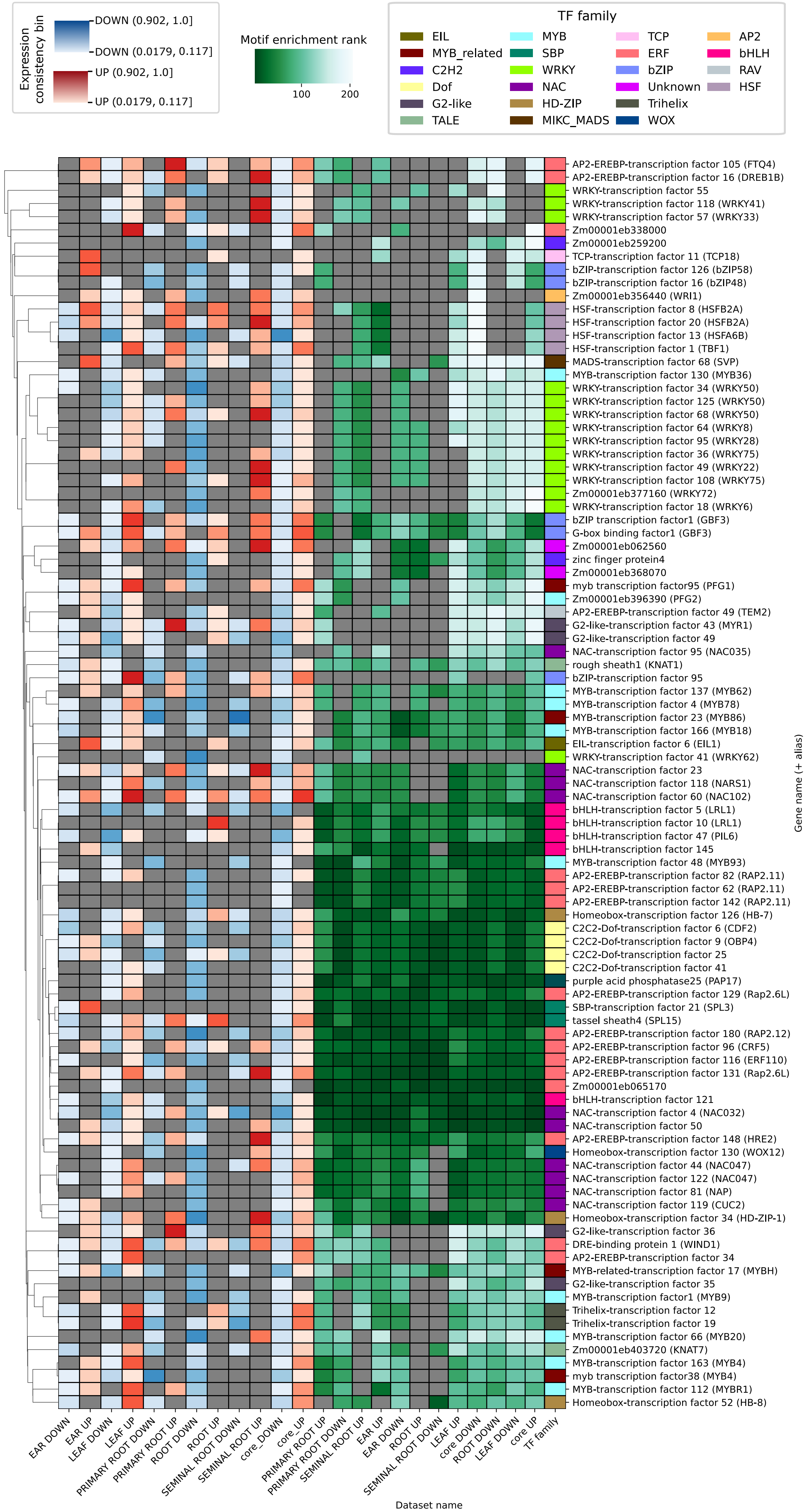


**Supplementary Figure 7. Complete clustered heatmap showing the motif enrichment rank of differentially expressed transcription factors (TFs) in the inferred gene regulatory networks (GRNs).** TFs (y-axis) are shown for the “all iCREs”-based GRNs, inferred using sets of iCREs that are associated with tissue-specific (x-axis) up-regulated genes in response to drought. The row annotations correspond to the activation and repression consistency score bins and the TF family.

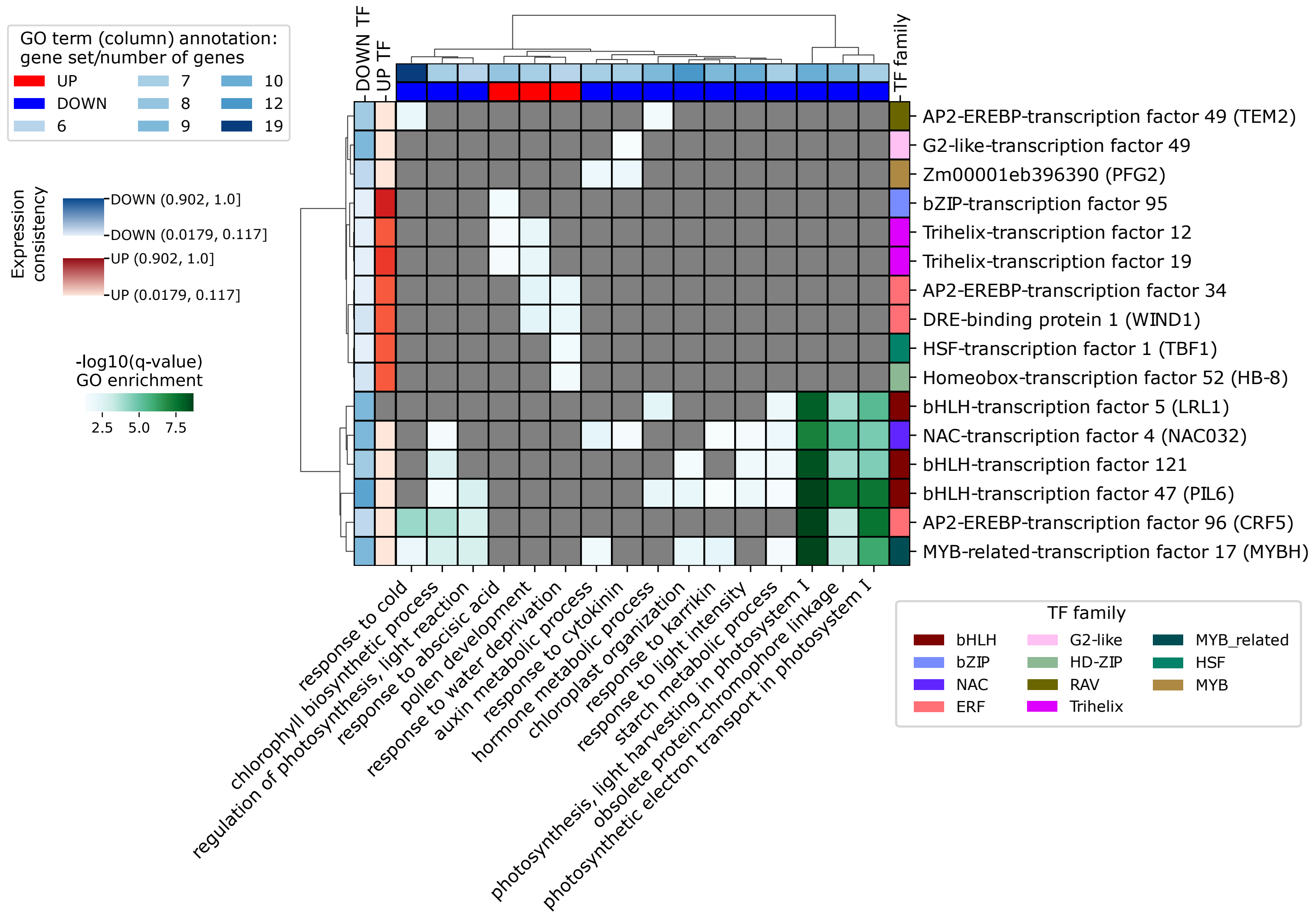


**Supplementary Figure 8. Clustered heatmap of gene ontology (GO) enrichment for target genes of predicted regulators in the inferred drought-responsive gene regulatory networks.** The -log10(q-value) of GO (x-axis) enrichments are displayed among the predicted target genes of up- and down-regulated transcription factors (TFs) in response to drought in leaf (y-axis). The row annotations indicate the activation and repression consistency score of the TFs, and the TF family. The column annotation corresponds to the gene set (up- or down-regulated marker genes in response to drought in leaf) that yielded the indicated GO enrichment, and the number of target genes in that GO category. For visualization purposes, the enriched GO terms with 5 or fewer target genes were discarded, as well as TFs with 3 or less GO terms enriched

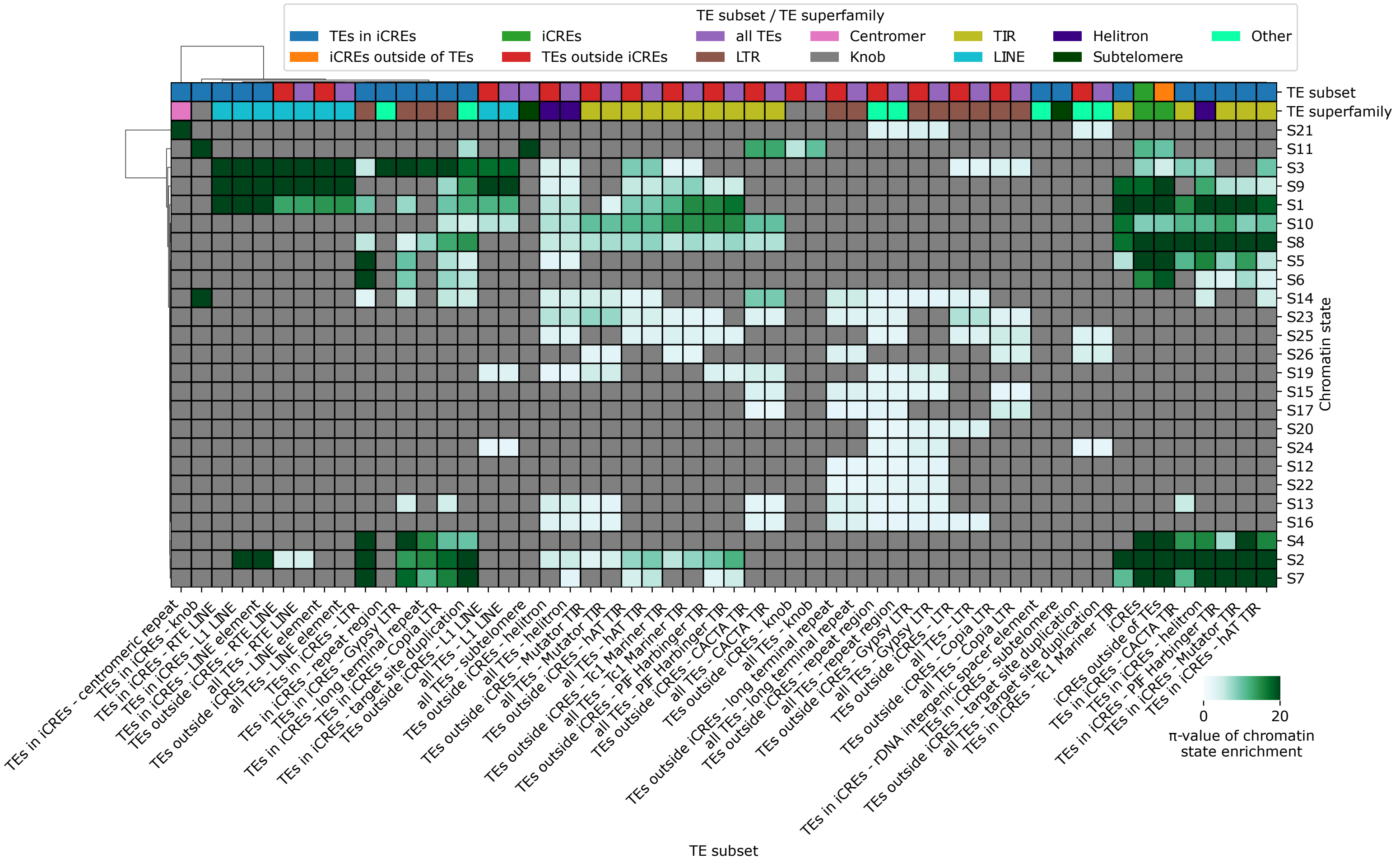


**Supplementary Figure 9. Chromatin states (CS) enrichment per TE superfamily and repetitive elements type.** Clustered heatmap showing the CS (y-axis) enrichment π-value of different TE subsets split per TE superfamily or repetitive elements type (x-axis). The columns are annotated based on the TE superfamily/repetitive elements type, and based on the TE subset (all TEs, TEs within iCREs, and TEs outside iCREs).

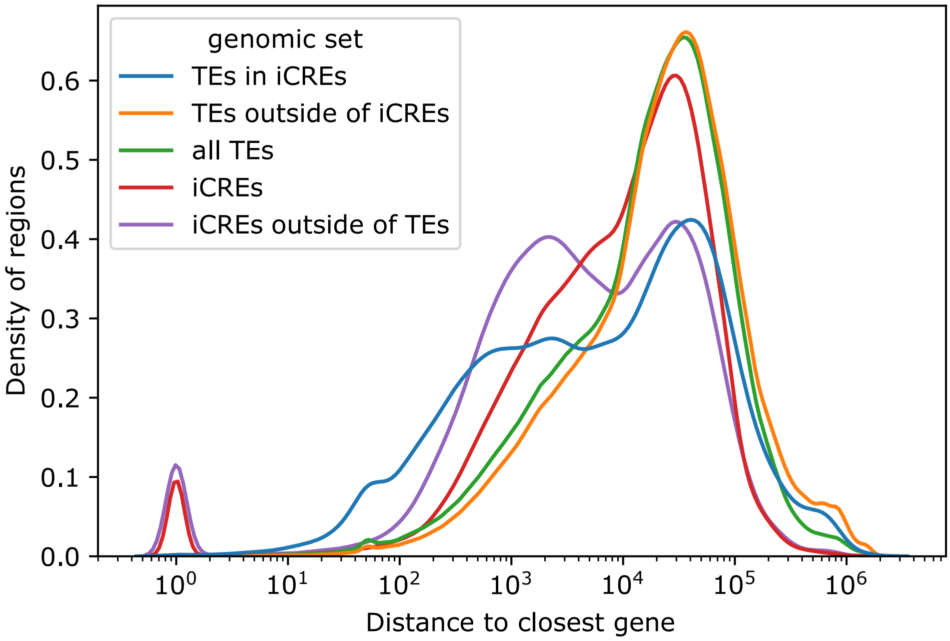


**Supplementary Figure 10. Distribution of the distance to the closest gene for the indicated sets of genomic regions.**
